# Real-time spatial evolution of the fMRI response to photobiomodulation in the healthy human brain

**DOI:** 10.1101/2025.11.17.688884

**Authors:** Joanna Chen, Hannah Van Lankveld, Xiaole Z. Zhong, J. Jean Chen

## Abstract

Photobiomodulation (PBM) is a non-invasive therapeutic technique that uses low-level near-infrared light to influence mitochondrial metabolism and stimulate neural function. While PBM is increasingly used to improve cognitive and clinical outcomes, its *in vivo* physiological mechanisms in humans remain poorly characterized. In this study, we used BOLD-fMRI to investigate and quantify the temporal and spatial dynamics of transcranial PBM-induced brain activity in young, healthy adults, while incorporating multiple stimulation parameters (wavelength, irradiance, frequency) and transcranial delivery sites (right forehead and intranasal). A time-lagged correlation analysis revealed distinct spatiotemporal patterns of positive and negative BOLD responses that evolved over tens of seconds across both cortical surface and subcortical regions during stimulation. Notably, these effects propagated across brain regions that are potentially mediated by functional networks, were dose-dependent, and were modulated by individual skin tone. This work provides the first real-time, whole-brain mapping of PBM-induced hemodynamic changes in humans, offering new insights into dose-response characteristics and delivery-specific dynamics underlying PBM neurophysiology.

## Introduction

Photobiomodulation (PBM) is a non-invasive neuromodulation technique that involves the application of low-intensity light in the red to near-infrared (NIR) spectrum to stimulate and enhance the function of neural tissue, cellular processes, and tissue repair (Hamblin 2016). NIR light, able to penetrate human skin and tissue to sufficient depth to effectively stimulate tissue at targeted regions (Wu et al. 2020; Stepanov et al. 2022; Zivin et al. 2009; Nizamutdinov et al. 2021), interacts with various chromophores and metabolites in the human body to induce beneficial physiological responses. Specifically, PBM is able to increase the bioavailability of adenosine triphosphate (ATP) by facilitating its production through the absorption of light by cytochrome c oxidase, the terminal enzyme of the electron transport chain (Srinivasan et al. 2013; Cardoso et al. 2022; Hamblin 2018). Located in the membrane of mitochondria, CCO facilitates the reduction of oxygen to water in cellular respiration, which enables the production of ATP in the process of oxidative phosphorylation. Additionally, PBM also induces the dissociation of bound inhibitory nitric oxide (NO) from CCO, which further drives ATP production while also creating a vasodilatory effect (Yokomizo et al. 2022; Quirk and Whelan 2020; Brown 2000; Kashiwagi et al. 2024). This ultimately leads to an increase in cerebral blood flow (CBF), neuronal activity, and metabolic activity in stimulated areas in PBM.

PBM is currently in several clinical trials for the treatment of a variety of neurological conditions, such as stroke, Parkinson’s disease, Alzheimer’s disease, and depression (Montazeri et al. 2021). Recent studies have demonstrated the various therapeutic benefits that PBM can elicit; for example, experimentally, it has been found that PBM can improve symptom management and outcomes in Alzheimer’s-related dementia (Nizamutdinov et al. 2021). Additionally, evidence also suggests that PBM can enhance neurocognitive functions in healthy humans as well (Gonzalez-Lima 2021; Zhao et al. 2022). However, while the photo-activity of CCO and the clinical benefits of PBM have been well-characterized through cell culture, *in vitro*, and clinical cognitive testing, there is still limited understanding of the precise *in vivo* physiological mechanisms of the PBM response in humans. In order to add onto existing knowledge of the action of these mechanisms *in vivo*, in recent years, there have been several studies focusing on quantifying the physiological response to PBM using BOLD-fMRI.

Compared to other neuroimaging modalities, like electroencephalography (EEG) and functional NIR spectroscopy (fNIRS), BOLD-fMRI offers global coverage of the brain while providing excellent temporal and spatial resolution (Tong 2022). Several PBM-fMRI studies have shown that PBM can increase functional connectivity in the Alzheimer’s population and chronic stroke patients (Chao 2019a; Naeser et al. 2020; Rashidi-Ranjbar et al. 2025). Additional studies examining Alzheimer’s have also reported decreases in fractional amplitude of low-frequency fluctuations as a result of PBM (Gaggi et al. 2024). PBM-fMRI studies in young, healthy adults have also found increased functional connectivity during PBM stimulus, yet no significant difference in functional connectivity when comparing pre- and post-stimulus effects (El Khoury, Mitrofanis, and Henderson 2019; Khoury, Mitrofanis, and Henderson 2021; Dmochowski et al. 2020). This raises interesting questions towards changes in brain circuitry and the active physiological response during response versus after response, as well as investigating the separation of the online and offline effects of PBM as separate mechanisms. However, the use of pre- and post-comparison analysis, a method that dominates PBM BOLD-fMRI studies, has limited ability in the characterization and interpretation of how the brain dynamically responds to the start of PBM stimulus, as well as when the stimulus ends.

As such, the actual mechanisms of action in the *in vivo* response to light remain poorly understood. Furthermore, the transmission of the PBM response, which has been reported as more global rather than focal (Dole et al. 2023), from the original site of stimulation also remains unclear. Applying dynamic analysis to the *in vivo* BOLD-fMRI response to PBM could provide valuable insight into the interactions of these mechanisms with light, as well as a better characterization of how the response begins, spreads, and ends. Additionally, biophysical modeling in our work and others predicts response dependence on wavelength, irradiance, skin tone, and, in the case of pulsed PBM, pulsation frequency, which all remain untested *in vivo* (Van Lankveld et al. 2024) – dynamic analysis could discern the specific effects of dose-dependency and site-specificity on the online and offline response.

In this study, we aim to characterize the evolution and spread of the brain’s physiological response during and after PBM in healthy adults, using varied doses of light delivery and stimulation sites, as well as quantifying demographic variations like skin tone. We hypothesize that variations in stimulation parameters and skin tone would elicit differences in the spatial and temporal evolution of BOLD-fMRI responses. We also hypothesize that PBM administered at different stimulation sites would result in regional differences and spatial localization of response.

## Methods

### Subject recruitment and ethical approval

A total of 47 healthy young adults were recruited through the Baycrest Participants Database, consisting of individuals from Baycrest and local communities. Participants were screened prior to participation to ensure there was no history of neurological or physiological disorders, malignant disease, or the use of medications or supplements that could have influenced the study. All experiments were conducted in accordance with the Baycrest Research Ethics Board (REB) guidelines, and all participants provided written informed consent, approved by the REB. Over the course of the active study, 2 participants withdrew early, one due to claustrophobia and one due to technical calibration issues, resulting in a final participant pool of 45 subjects (23 M, mean age = 26.0 ± 3.4 years / 22 F, mean age = 23.0 ± 2.8 years).

### Skin tone assessment

To account for individual differences in skin optical properties that may affect light penetration during PBM, we targeted an equal representation of three skin colour categories (light, intermediate, dark) during recruitment. Individual skin colour was quantified by measuring forehead skin pigmentation and melanin level using a CM-600D spectrophotometer and calculating the individual typological angle (ITA) (Konica Minolta, Tokyo, Japan). Participants were then organized into three groups (Groups 1, 2, 3) according to their measured ITA, with Group 1 corresponding to the lightest skin and Group 3 the darkest. For additional details about skin tone quantification and ITA calculation, additional information is attached in the Supplementary Materials.

### MRI Acquisition and Scanning

fMRI data were acquired using a Siemens Prisma 3 Tesla System (Siemens, Erlangen, Germany), which employed a 20-channel phased-array head coil reception and body coil transmission. T1-weighted 3D anatomical images (sagittal, 234 slices, 0.7 mm isotropic resolution, TE = 2.9 ms, TR = 2240 ms, TI = 1130 ms, flip angle = 10°) were acquired for positioning. We acquired BOLD-fMRI data with dual-echo (DE) pseudo-continuous arterial spin labelling (pCASL) (courtesy of Danny J. J. Wang, University of Southern California) for recording BOLD dynamics (TR = 4.5 s, TE1 = 9.8 ms, TE1 = 30 ms, post-labelling delay = 1.5 s, labelling duration = 1.5 s, flip angle = 90°, 3.5 mm isotropic resolution, 35 slices, slices gap = 25%, scanning time = 4 minutes). Participants were scanned while viewing naturalistic landscapes and imagery to reduce random mind-wandering as well as ensure their eyes remained open, thereby improving the reproducibility of the results.

To test for the existence of brain temperature increases during PBM, MR thermometry data were collected for the sessions corresponding to the 1064nm wavelength delivered at the highest irradiance (200 mW/cm²) at both 10 Hz and 40 Hz (N = 30 participants). For the full details regarding MR thermometry, data analysis, and results, see Supplementary Materials for more information.

### Laser Instrumentation and protocol design

Based on existing PBM literature, the stimulation parameters chosen for this study consisted of two wavelengths (808 nm and 1064 nm), two pulsation frequencies (10 Hz and 40 Hz), and three optical power densities for each positioning (**Table 1**). The target sites where light was to be delivered were chosen to be to the right prefrontal cortex for forehead positioning (tPBM) and at the left nostril for nasal positioning (iPBM) (**Fig. 1a**). To characterize subject variability in anatomical differences in nasal passage, the nasal laser distance to the brain was measured using Freeview’s built-in measure tool, as seen in **Fig. 1b**. Two MR-compatible Class 3 laser systems were acquired from Vielight Inc: MDL-III-808-1W and MDL-III-1064-1W, which delivered 808nm and 1064nm light respectively. The systems were connected to a waveform generator to produce the desired duty cycle (50%), and light was delivered through 10-meter, 400 μm optic cables fitted to a custom headpiece from the system to the MRI. This allowed the laser systems to remain external to the MRI environment while ensuring remote controllability of the stimulation parameters, preventing potential interference. This also meant that the participant remained blinded to the stimulation paradigm to avoid any placebo effects. Two vitamin E capsules were also placed to serve as fiducial markers to pinpoint the location of stimulation in the anatomical and DE-pCASL images.

**Figure 1:**
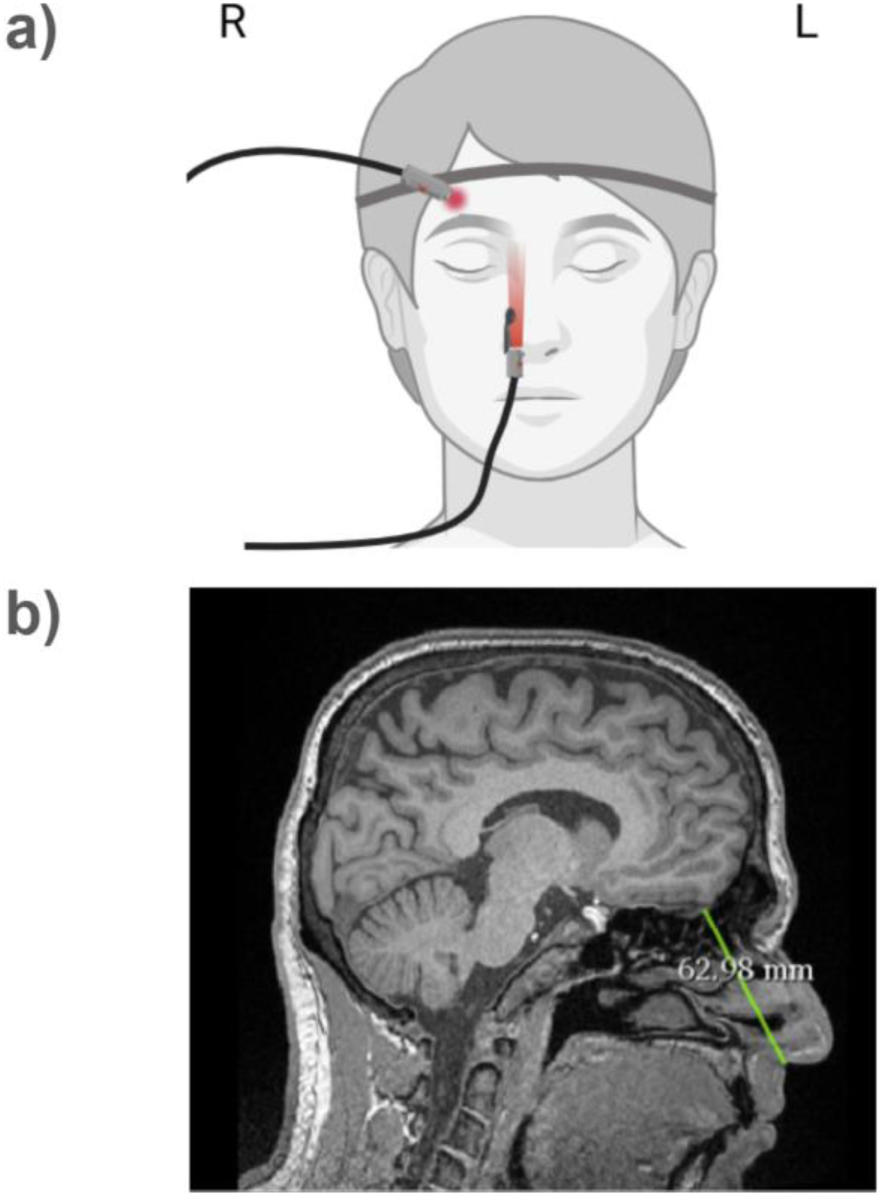
Forehead and intranasal laser positioning. **a) Laser positioning during stimulus:** The forehead laser was secured with a band to the forehead and positioned perpendicular to the right prefrontal cortex. The intranasal laser was secured with a nostril clamp at the right nostril. As both lasers were secured during the entire scans, participants were blinded to when the laser was on or off. **b) Measurement of intranasal laser distance to the brain:** This was calculated manually through Freeview’s built-in measure tool, using the subnasale (the intersection of the philtrum and base of the nose), angle of nasal passage, and grey matter as landmarks in the right nostril passage.

**Table 1.** PBM protocol group design.

| Scan # | Site | Protocol 1 (P1) | Protocol 2 (P2) | Protocol 3 (P3) |
| --- | --- | --- | --- | --- |
| 0 |  | Anatomical | Anatomical | Anatomical |
| 1 | tPBM | 1064 nm, 100 mW/cm<sup>2</sup>, 10 Hz | 1064 nm, 150 mW/cm<sup>2</sup>, 10 Hz | 1064 nm, 150 mW/cm<sup>2</sup>, 40 Hz |
| 2 | iPBM | 808 nm, 9 mW/cm<sup>2</sup>, 10 Hz | 808 nm, 5 mW/cm<sup>2</sup>, 10 Hz | 808 nm, 5 mW/cm<sup>2</sup>, 40 Hz |
| 3 | tPBM | 808 nm, 100 mW/cm<sup>2</sup>, 10 Hz | 808 nm, 150 mW/cm<sup>2</sup>, 10 Hz | 808 nm, 150 mW/cm<sup>2</sup>, 40 Hz |
| 4 | iPBM | 1064 nm, 9 mW/cm<sup>2</sup>, 10 Hz | 1064 nm, 5 mW/cm<sup>2</sup>, 40 Hz | 1064 nm, 5 mW/cm<sup>2</sup>, 40 Hz |
| 5 | tPBM | 1064 nm, 200 mW/cm<sup>2</sup>, 10 Hz | 1064 nm, 200 mW/cm<sup>2</sup>, 40 Hz | 1064 nm, 100 mW/cm<sup>2</sup>, 40 Hz |
| 6 | iPBM | 808 nm, 7 mW/cm<sup>2</sup>, 10 Hz | 808 nm, 7 mW/cm<sup>2</sup>, 40 Hz | 808 nm, 9 mW/cm<sup>2</sup>, 40 Hz |
| 7 | tPBM | 808 nm, 200 mW/cm<sup>2</sup>, 10 Hz | 808 nm, 200 mW/cm<sup>2</sup>, 40 Hz | 808 nm, 100 mW/cm<sup>2</sup>, 40 Hz |
| 8 | iPBM | 1064 nm, 7 mW/cm<sup>2</sup>, 10 Hz | 1064 nm, 7 mW/cm<sup>2</sup>, 40 Hz | 1064 nm, 9 mW/cm<sup>2</sup>, 40 Hz |
| **M/F** |  | 8 M / 8 F | 7 M / 7 F | 9 M / 7 F |

Each participant was assigned to one of three protocol design groups, with each protocol composed of eight unique combinations of stimulation parameters (**Table 1**). The protocols varied between the three optical power densities, two wavelengths, and two pulsation frequencies for each positioning in order to thoroughly evaluate effects and potential interactions between PBM parameters on brain activity. Each participant completed eight separate fMRI scans, with each scan being associated with a specific parameter setting. Equal distribution of ITA grouping and sex was ensured during protocol assignment (**Table 2**).

**Table 2.** Participant distribution by protocol and melanin group.

|  | Protocol 1 | Protocol 2 | Protocol 3 |
| --- | --- | --- | --- |
| **Melanin Group 1** | 5 | 5 | 4 |
| **Melanin Group 2** | 6 | 5 | 6 |
| **Melanin Group 3** | 5 | 4 | 6 |
| **# of Subjects** | 14 | 16 | 15 |

Each fMRI scan was 12 minutes long and followed a block design (OFF-ON-OFF) stimulation paradigm (see **Fig. 2a**): 4 minutes of baseline, 4 minutes of PBM stimulation (DURING STIMULUS), and 4 minutes of post-stimulation (POST-STIMULUS). This duration was chosen as it is within the range used in PBM literature and allows us to monitor both online and offline effects (Wang et al. 2019; Dmochowski et al. 2020). Combining the 4-minute baseline period, the 4-minute post-stimulation period, and the preparation period between consecutive scans, there was a minimum of 20 minutes of rest between periods of active PBM stimulation (see **Fig. 2b**).

**Figure 2:**
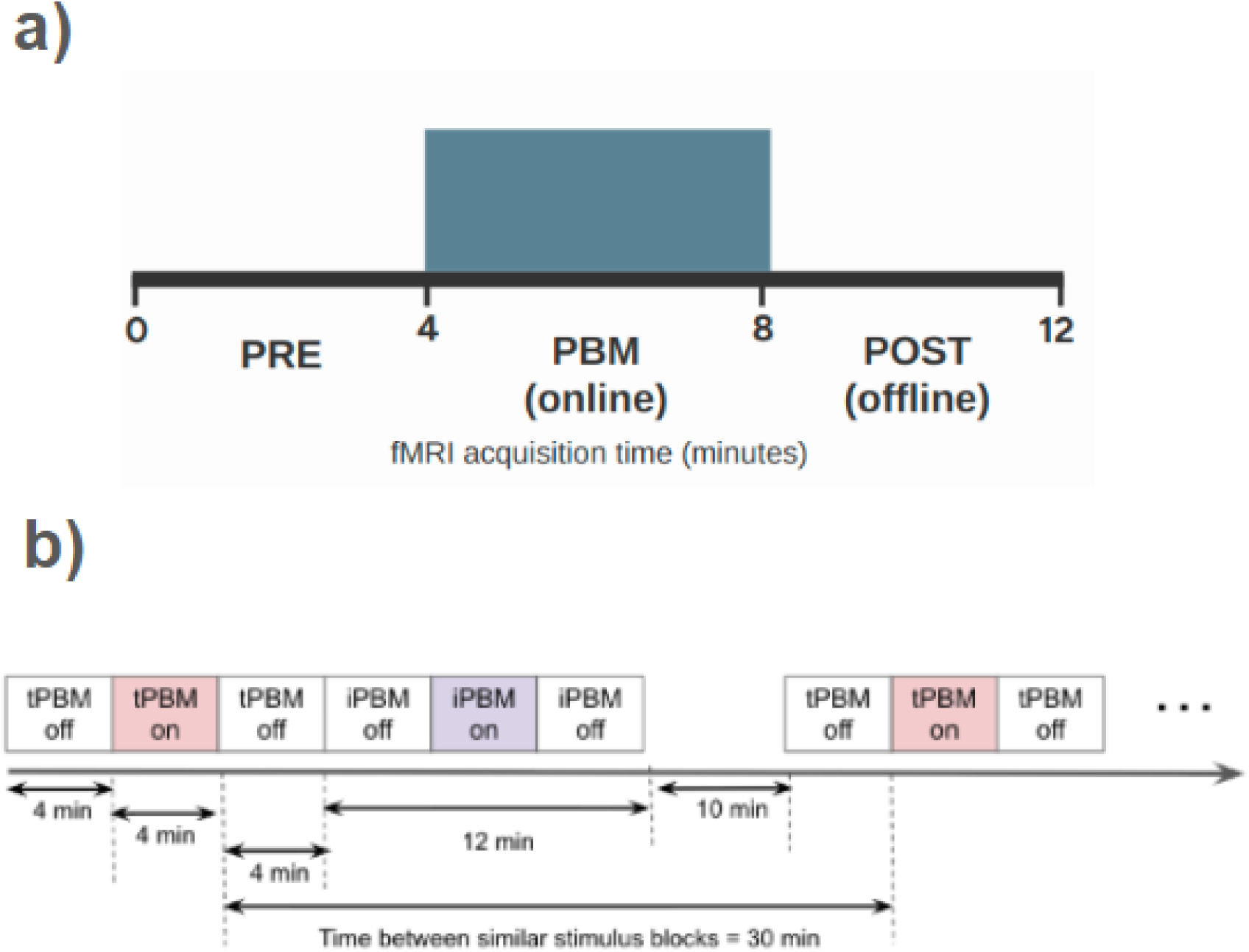
a) PBM stimulation paradigm and simultaneous fMRI DE-pCASL recording. The shaded region represents the time frame in which the laser was turned on. The same timing is used for both tPBM and iPBM stimulations. **b) Timing of laser on/off periods between consecutive scans.** tPBM and iPBM were delivered in alternation, such that there were 20 minutes of separation between stimulation at the same site.

### Data analysis

#### fMRI preprocessing

Preprocessing was performed using the FMRIB Software Library (FSL) (Analysis Group, FMRIB, Oxford, UK). The BOLD signal was calculated by surround-averaging images from the second echo of DE-pCASL data. Steps during the preprocessing stage included brain extraction, motion correction, distortion correction, slice timing, spatial transformation into Montreal Neurological Institute (MNI152) space, thresholding, and temporal and spatial outlier removal. Additional steps included masking out ventricles, the application of a bandpass filter to the 0.0005-0.006Hz range (determined based on stimulus timing), and normalization. The data was normalized using the following formula:

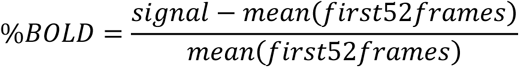

#### Time-lagged correlation analysis for spatial evolution

To apply correlation analysis of the predicted response to different lag times on a global scale, we used the Rapidtide (v2.9.6) package, written in Python (Frederick, B, 2024). Rapidtide implements an algorithm called RIPTiDe (Regressor Interpolation at Progressive Time Delays) that performs rapid time delay analysis in filtered fMRI data to find lagged correlations between each voxel’s time-series and a reference time-series, referred to as a “probe regressor”. We chose to use the Rapidtide toolbox to estimate the PBM response in fMRI data because: (i) it is a citable open-source software package demonstrated to be successful in providing exquisite information on the spatial dynamics in the BOLD response (Tong et al., 2011; Erdogan et al., 2016; Champagne et al., 2022) (ii) with a data-driven probe regressor, we do not have to assume a hemodynamic response function for PBM, which is unknown. In our analysis, we used the initial stimulus design (see **Fig. 3a**) as a positive probe regressor and inverted the design to create a negative probe regressor. As such, our analysis was set up to assess explicit positive and negative responses (**Fig. 3a**). For each subject, Rapidtide produced a whole-brain maximum-correlation map with the regressor. This map was then thresholded for significance (p<0.01). The thresholded Rapidtide map was then first used to generate a map of significant lag times to the regressor by using MATLAB’s xcorr function with the 4D preprocessed data. The lag times of the shifted copies were then used to threshold the original map into submaps of time intervals of 4.5 s, which allowed us to depict the frame-by-frame evolution of the tPBM response (see **Fig. 3b**).

**Figure 3:**
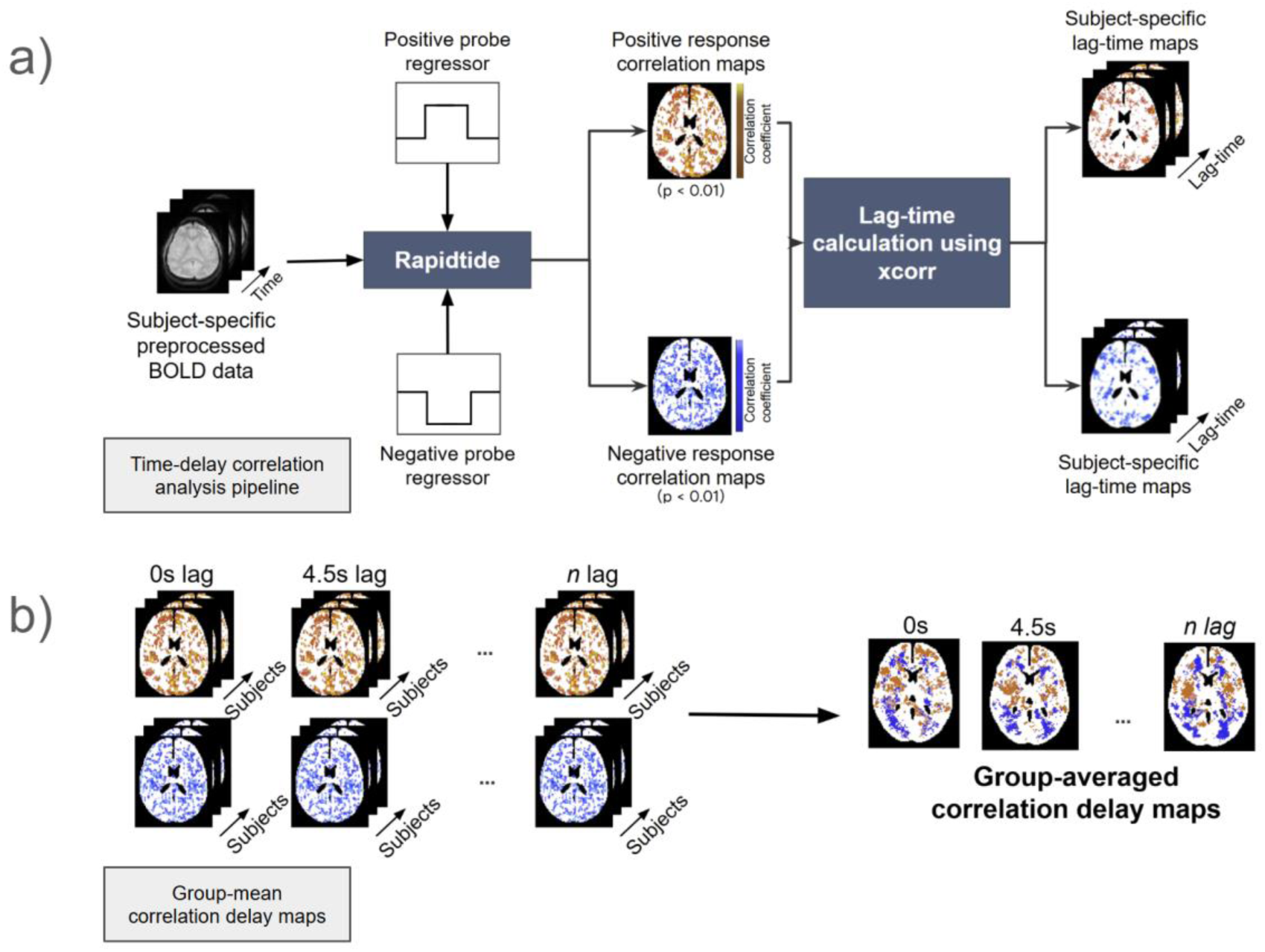
Preprocessing and analysis pipeline. **a)** Time-delay correlation analysis: Correlation maps from each subject and PBM session were generated from significant voxels correlating to the two regressor probes (p<0.01) in Rapidtide. These maps were then used for cross-correlation analysis using MATLAB. **b)** Group-mean correlation delay maps: generated by segmenting voxels based on lag-time, then averaging across the group and thresholding for 95th percentile commonality.

#### Quantification of the temporal evolution

After performing FDR-correction using the Benjamini-Hochberg method on the thresholded Rapidtide maps to correct for multiple comparisons (Benjamini and Hochberg 1995), surviving voxels with a corrected p-value<0.05 were then binarized to form a mask, which was then used to extract the average %BOLD timeseries for each scan of each subject. From each %BOLD time series extracted from averaging across all significant voxels in each fMRI scan, different aspects of the temporal evolution of the response were quantified and submitted to linear mixed-effects analyses (LMEs) (see **Figure 4a**). For all parameters, outliers were removed prior to LME analysis. For the time parameters, a 10-second sliding window was used to average the time course to ensure better-fitting exponential curves.

**Figure 4:**
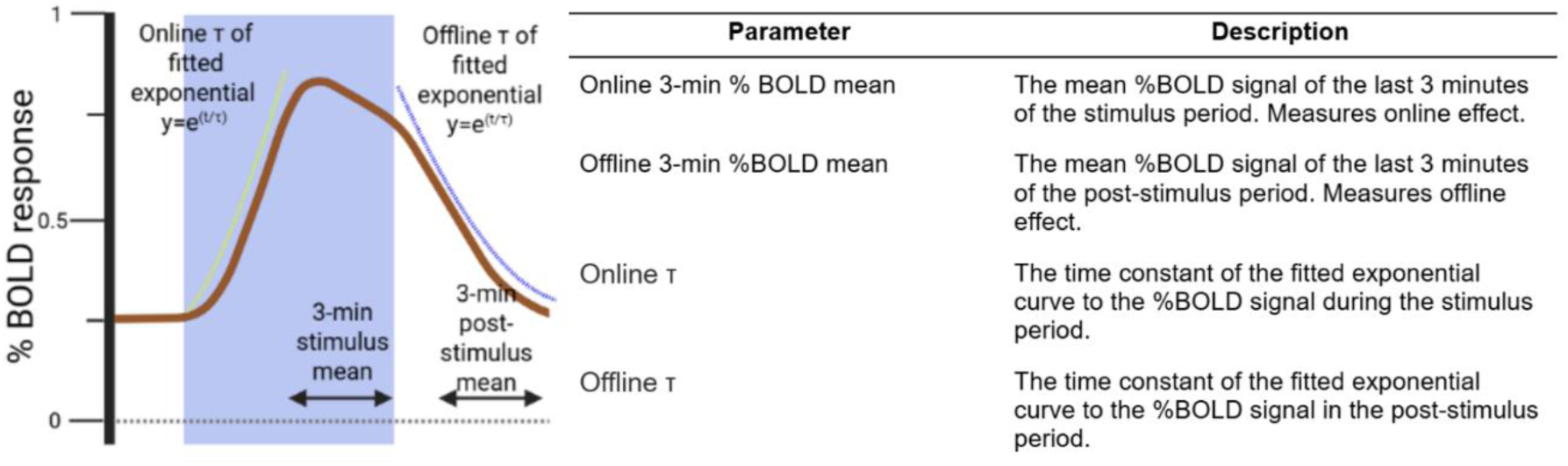
Linear mixed effects parameters: a) Temporal response BOLD parameters. Parameters were extracted from subject-specific %BOLD time series as illustrated here and then submitted to linear mixed effects analysis to assess group-level effects.

#### Linear Mixed Effects analysis

All parameters were submitted to a custom two-stage Linear Mixed Effects (LME) modeling pipeline implemented in MATLAB (Natick, MA, USA). The temporal-response parameters were each in turn tested as the dependent variable. Independent variables considered for fixed effects included wavelength, irradiance, pulsation frequency, and ITA, as well as their interactions. With the exception of the ITA and intranasal laser distance, all fixed-effect variables are considered categorical. The subject ID was included as the random-effect variable.

In the first stage, we performed stepwise linear regression using stepwiselm in Matlab (Natick, MA, USA) to rule out predictors without significant contributions to the response variable. That is, in stepwiselm, variables (and their interactions) were added iteratively from a constant-only model, and excluded if their effect size did not meet a p < 0.05 False Discovery Rate (FDR)-corrected threshold using the Benjamini-Hochberg method (Benjamini and Hochberg 1995). This process was iterated using randomized ordering of dependent variables in stepwiselm to minimize order-related biases. In the second stage, only predictors that survived all iterations were then entered into the second stage LME model, where the response parameter variable was modelled, for example, as:

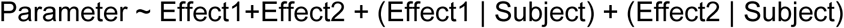

Where the parameter represents the BOLD response parameters (**Fig. 4b**) as the dependent variable, and Effect1, Effect 2, etc. were nominal fixed effects. The final statistical significance was evaluated again using FDR correction, with results considered significant if associated FDR-corrected p-values were lower than 0.05.

## Results

### MR thermometry

To evaluate potential temperature changes in the brain during tPBM, MR thermometry data were collected for the highest irradiance (i.e., at 1064 nm, 200 mW/cm², both 10 Hz and 40 Hz across 30 subjects). Group-level statistical comparison confirmed that even at the longest wavelength and highest irradiance, no measurable heating effects were produced in the targeted brain region. As the tPBM laser was positioned over the right forehead, aimed to stimulate the prefrontal cortex, this region was used as the target ROI for thermometry measurements. See Supplementary Materials for further details.

### The temporal BOLD response

The subject-specific %BOLD timeseries from each PBM session, extracted using subject-specific thresholded Rapidtide maps as masks (FDR-corrected p-value<0.05), were group-averaged to visualize the temporal BOLD response on a group level. In **Figure 5**, we summarize the group-averaged %BOLD response to different wavelength-frequency combinations for forehead (tPBM) and intranasal (iPBM) stimulation settings.

**Figure 5:**
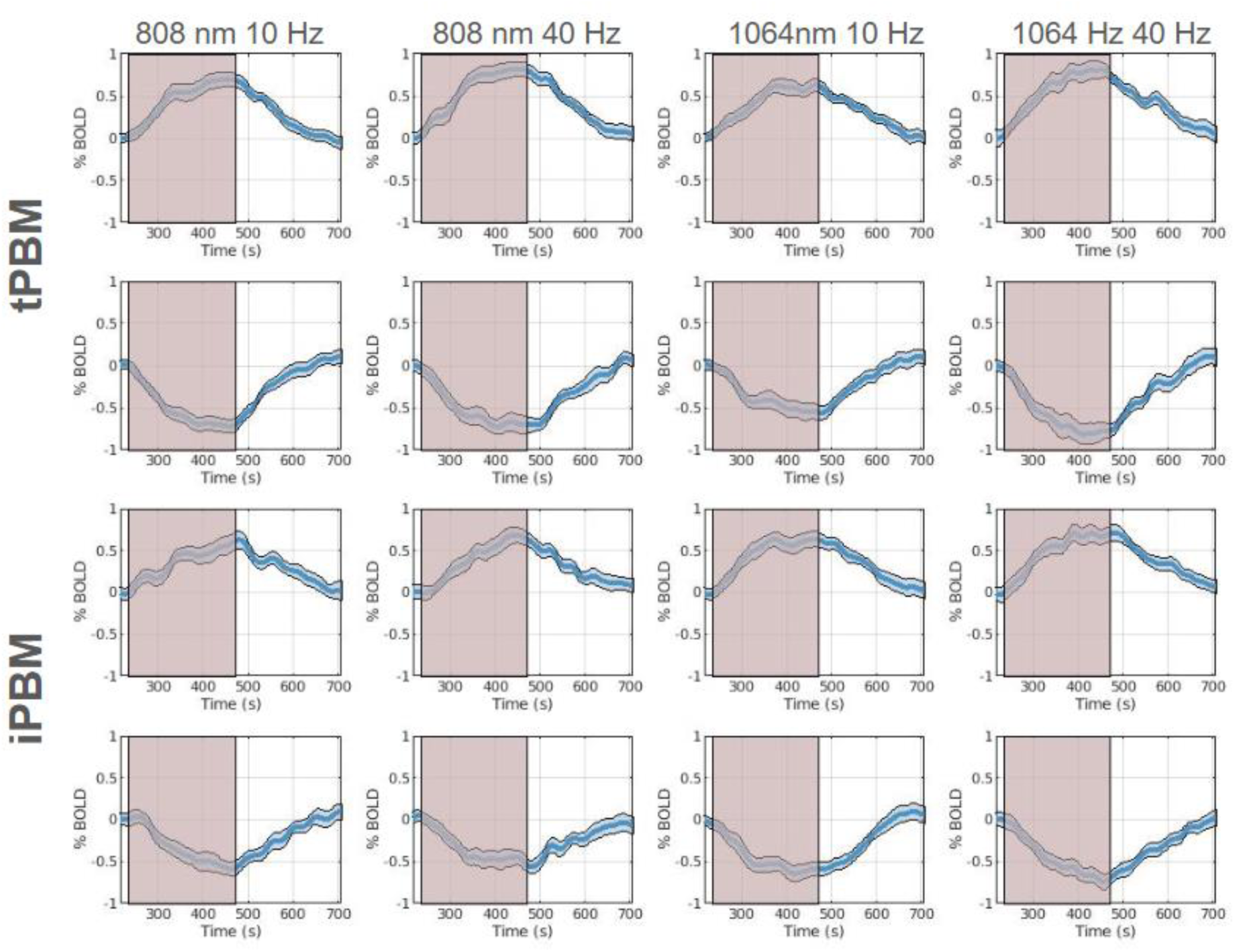
Group-averaged %BOLD timecourses. FDR correction was applied to maps following time-delay correlation analysis per subject and run, which were then used as masks to extract subject-specific %BOLD timecourses for each PBM session, averaged across significant voxels in that scan. The run-specific %BOLD time series was then averaged across protocol groups. The shaded red region represents the stimulus window in which the applied laser was on during the fMRI scan (progression shown through time axis). Blue shaded regions represent the mean standard error.

Interestingly, between both tPBM and iPBM, the magnitudes of the %BOLD response in both negative and positive response areas are similar, despite differences in irradiance levels used between the two stimulation sites. A distinct lagged response from the onset of stimulus can also be seen, with a steady rise or fall in %BOLD signal that begins to plateau on average about 150s after the stimulus begins.

### The cortical BOLD response

The group-averaged cortical surface maps depicting positive and negative BOLD responses for tPBM and iPBM are displayed in **Figures 6** and 7, respectively. Group-averaging was achieved by summing and averaging smoothed subject-specific correlation maps consisting of voxels whose maximum correlation coefficients survive the significance threshold (p<0.01) at each time lag. This group-averaged map was then thresholded at the 95^th^ percentile commonality to identify common regions of response across subjects and then mapped onto the FreeSurfer cortical surface.

**Figure 6:**
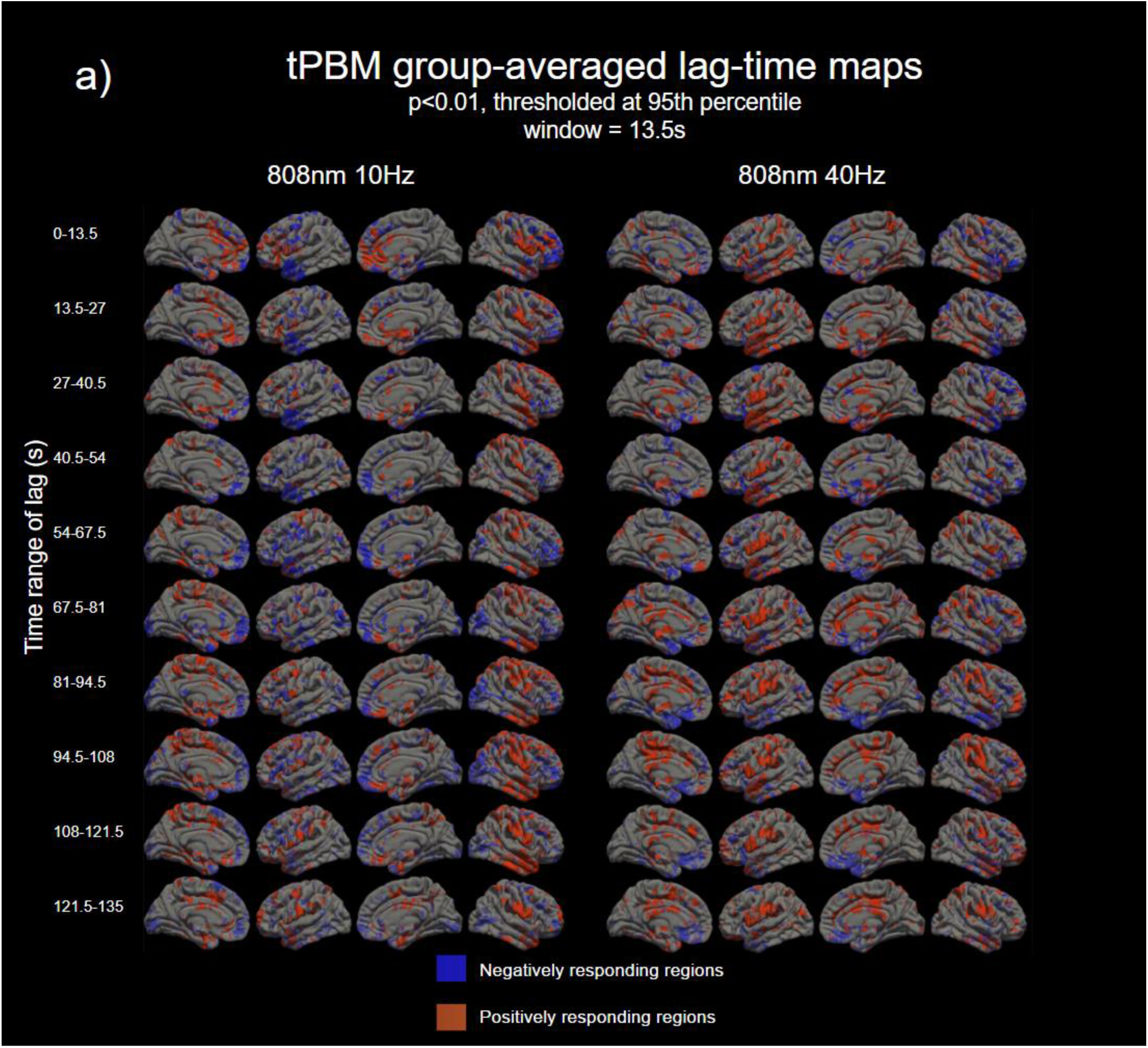
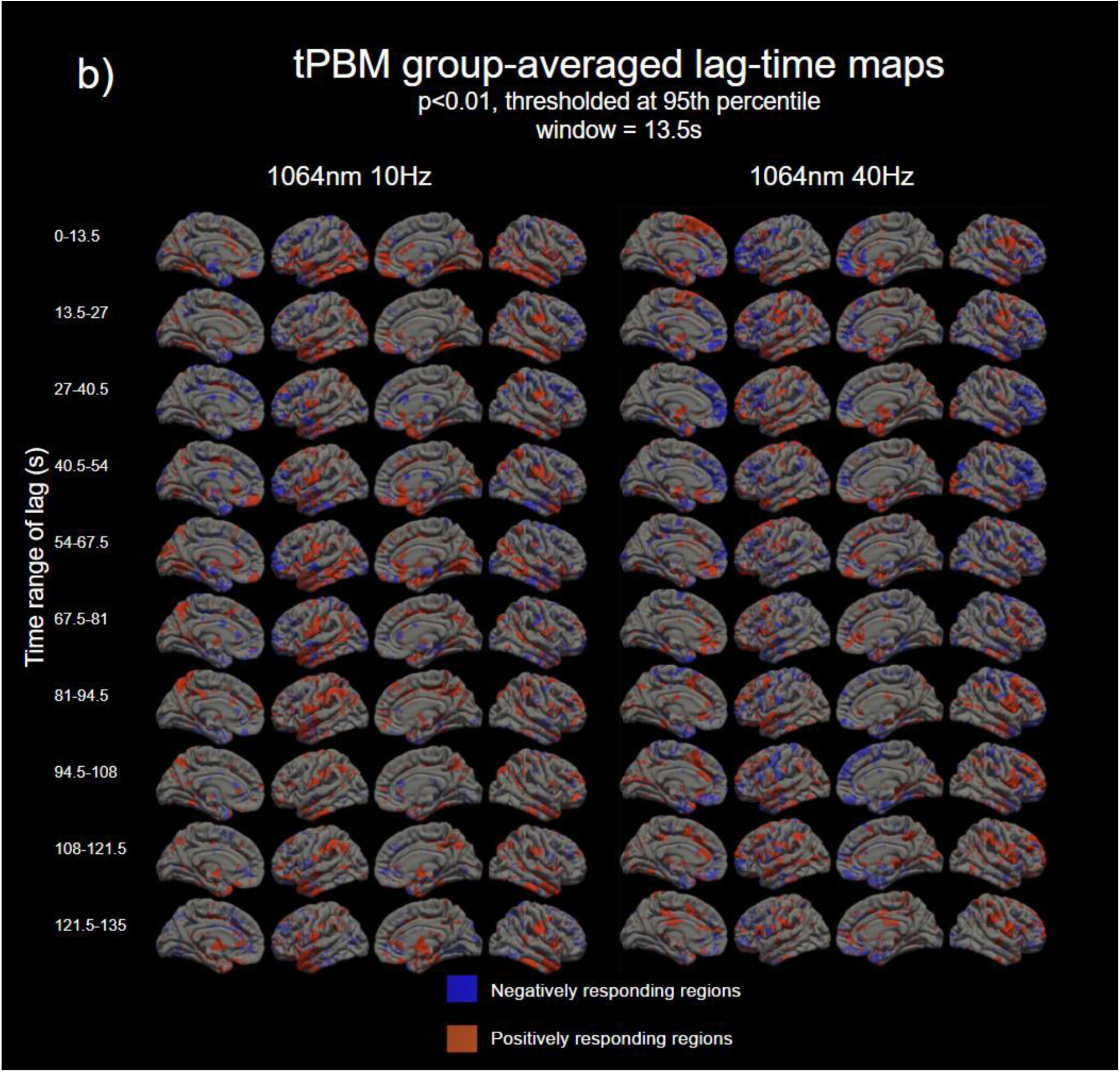
Group-averaged, window-by-window tPBM lag-time maps of **(a)** the 808 nm 10 Hz and 808 nm 40 Hz and **(b)** the 1064 nm 10 Hz and 1064 nm 40 Hz protocol combinations, mapped onto the cortical surface model, for the lateral and medial surfaces on both the left and right hemispheres. Blue and red correspond to regions where significant negative and positive BOLD responses were found (p<0.01). Maps were thresholded by the 95^th^ percentile of commonality to identify common regions of response.

**Figure 7:**
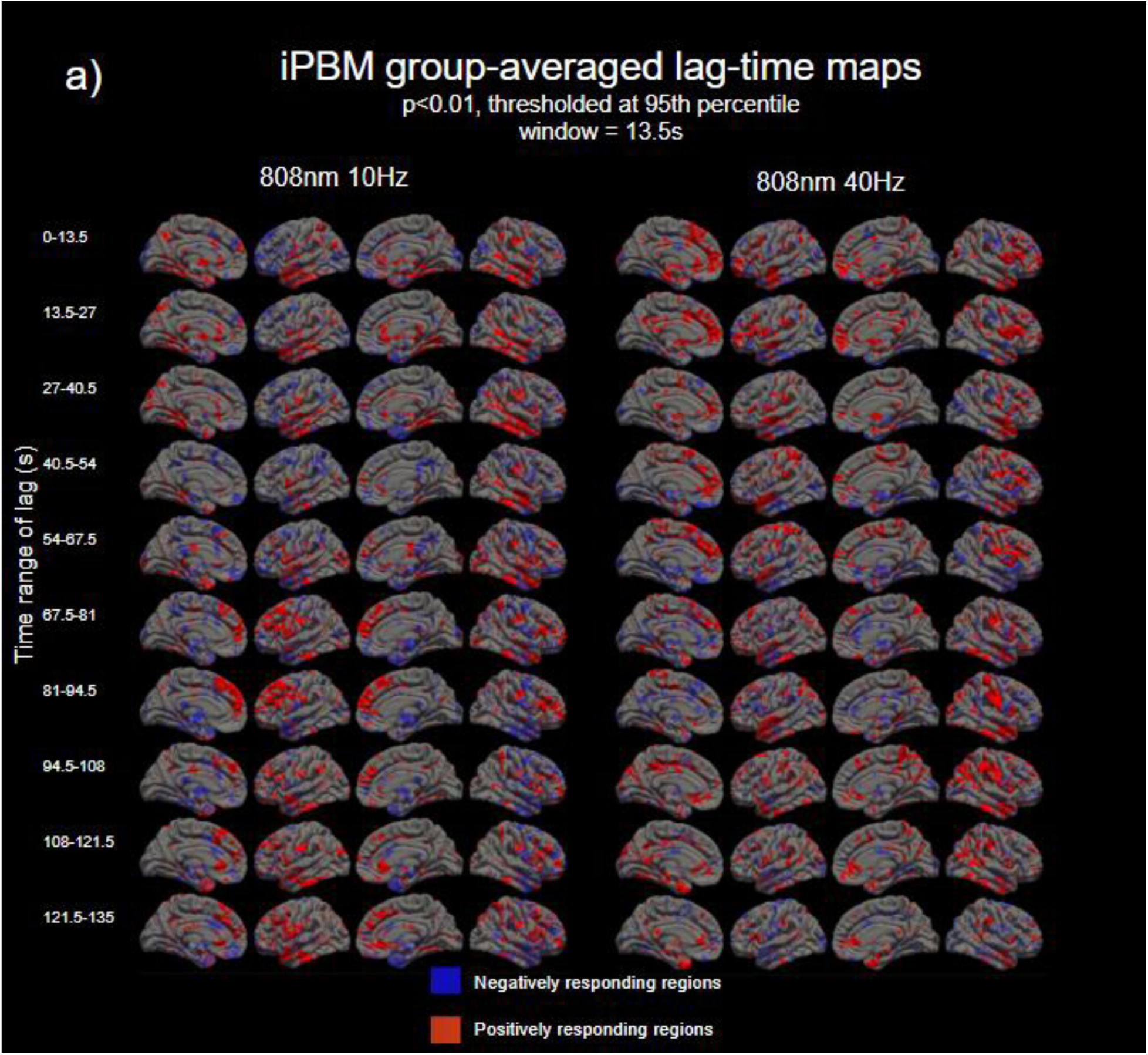
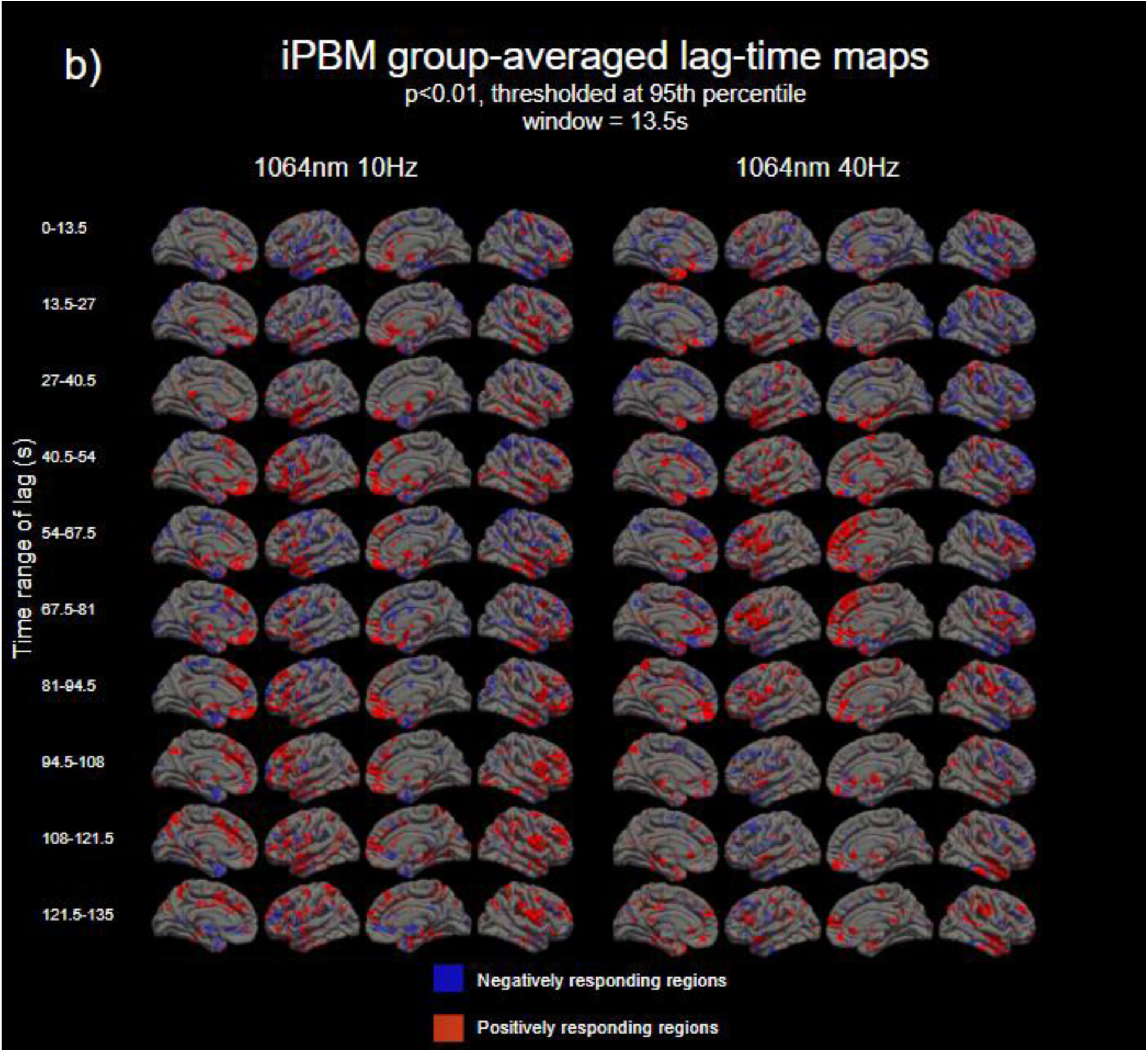
Group-averaged, window-by-window iPBM lag-time maps of **(a)** the 808 nm 10 Hz and 808 nm 40 Hz and **(b)** the 1064 nm 10 Hz and 1064 nm 40 Hz protocol combinations, mapped onto the cortical surface model, for the lateral and medial surfaces on both the left and right hemispheres. Blue and red correspond to regions where significant negative and positive BOLD responses were found (p<0.01). Maps were thresholded by the 95^th^ percentile of commonality to identify common regions of response.

In **Figures 6a** and **6b**, the time lags of the tPBM %BOLD response are displayed, organized by wavelength. For both 808 and 1064 nm stimulations, positive and negative BOLD responses can be seen throughout the cortical surface. Across most protocol groups, tPBM’s positive response tends to begin in the medial prefrontal cortex. Some positively responding areas can be seen on the lateral surface, but the lateral prefrontal cortex is largely dominated by negatively BOLD response regions. As time progresses, the positive response can be observed spreading either ventrally through the prefrontal cortex, or posteriorly, towards the back of the brain, passing briefly through the dorsomedial prefrontal cortex, superior frontal, paracentral lobule, and the precuneus. Following the precuneus activation, some positive response in the cingulate cortex can also be seen. This pattern is most strongly observed in 808 nm 10 Hz, 808 nm 40 Hz, and 1064 nm 40 Hz protocols.

In terms of characteristics of the negative BOLD response, 808 nm 10 Hz and 1064 nm 40 Hz demonstrate the most pronounced spatial extent for this type of response. However, the appearance of negative BOLD responses in 1064 nm 40 Hz is more concentrated in the first minute of stimulation (0-67.5s of lag time), while in 808 nm 10 Hz, negative responses continue to appear beyond this time frame, with delays as long as 135s. Additionally, the lagged negative responses in both 808 nm protocol groups tend to appear in the ventral medial prefrontal cortex, beginning 40s after the start of the stimulus. This pattern is not seen with the 1064 nm protocol groups, where the lagged response appears more superiorly as well as less extensive.

While the positive and negative BOLD responses appear to be symmetrical across both hemispheres in medial regions in all protocol groups, in the 808 nm protocol groups, some asymmetricality can be seen in the lateral regions. In 808 nm 10 Hz, the lateral temporal and parietal lobes in the left hemisphere tend to respond more negatively, while in the right hemisphere, these regions tend to respond positively. This trend is reversed in 808 nm 40 Hz; in the first minute of stimulus, the positive response tends to be concentrated in the left hemisphere, while the negative response is concentrated in the right hemisphere.

As seen in **Figures 7a** and **7b**, iPBM at 808 nm and 1064 nm elicits both positive and negative BOLD responses as well. As in tPBM, an initial positive response in the medial prefrontal cortex is also observed in iPBM, but it is much more ventral, likely due to the stimulation site. The medial response in iPBM can be seen strengthening and lingering in dorsal regions as time progresses, ranging from 20 - 90 s from the start of stimulus. This is a pattern of behaviour that is not seen in tPBM. Following the strengthening of the positive response in medial regions in iPBM, the positive response in lateral regions, such as the dorsolateral prefrontal area, as well as the temporal lobe, subsequently responds. Activation of the precuneus can also be observed around the same time, at around 81-94.5s in 808 nm 40 Hz, 1064 nm 10 Hz, and 1064 40 Hz.

The initial negative response in iPBM is the most pronounced in the 1064 nm protocol groups, with the 1064 nm 40 Hz negative response being more spatially extensive in the lateral hemisphere than the 1064 nm 10 Hz combination. Additionally, some asymmetry across hemispheres in the response can also be observed in 1064 nm 40 Hz, with the negative response dominating the lateral right hemisphere, while a positive response can be observed in the lateral left hemisphere.

### The subcortical response

Figures 8 and **9** depict group-averaged, subcortical maps of the evolution of the spatial response during stimulus. Like the cortical surface maps in Figs. 6 and 7, the subcortical maps were group-averaged by averaging the smoothed subject-specific correlation maps at each time interval (p<0.01). The group-averaged maps were then thresholded at 95^th^ percentile commonality and mapped onto the MNI152 brain.

**Figure 8:**
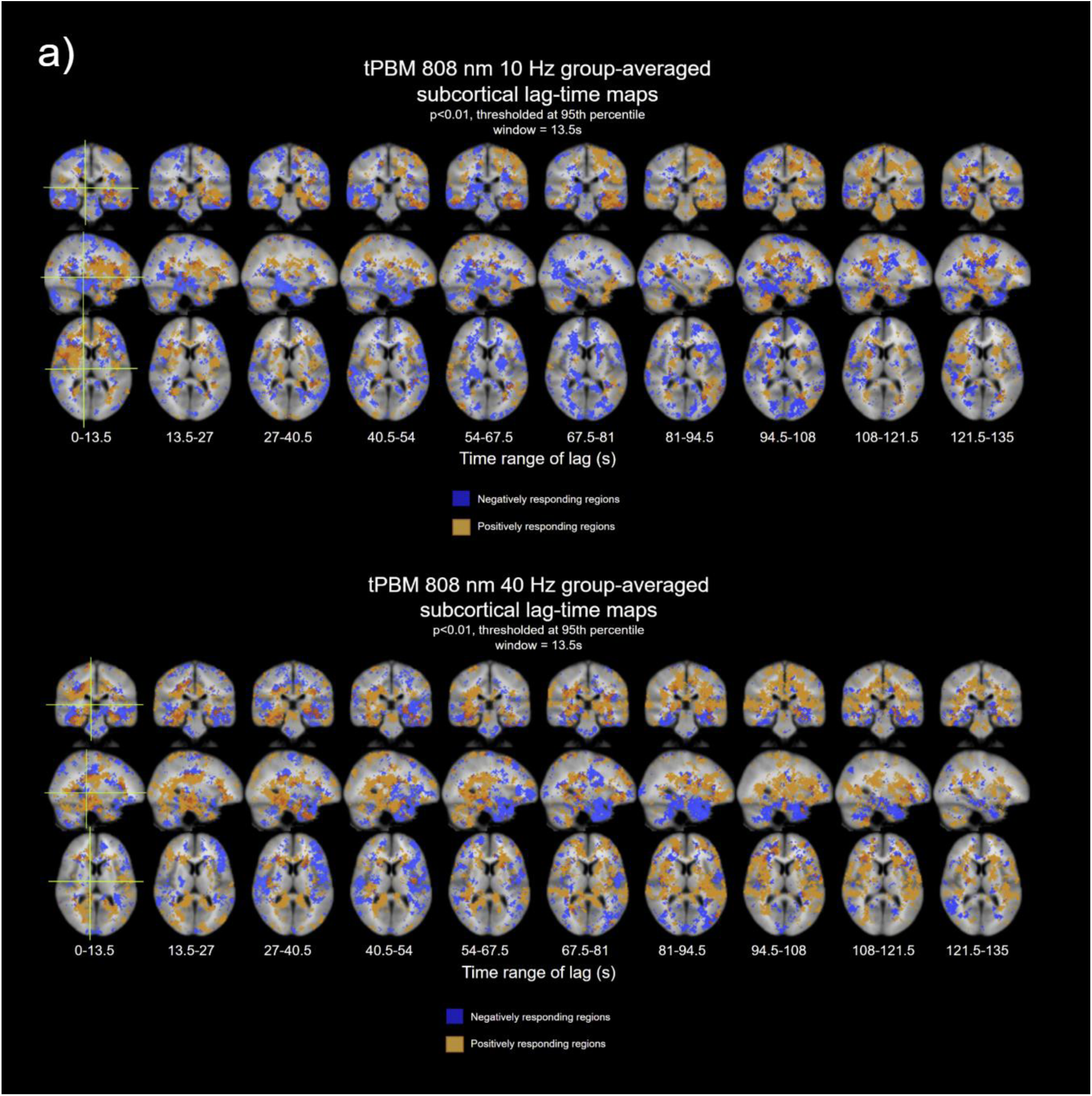
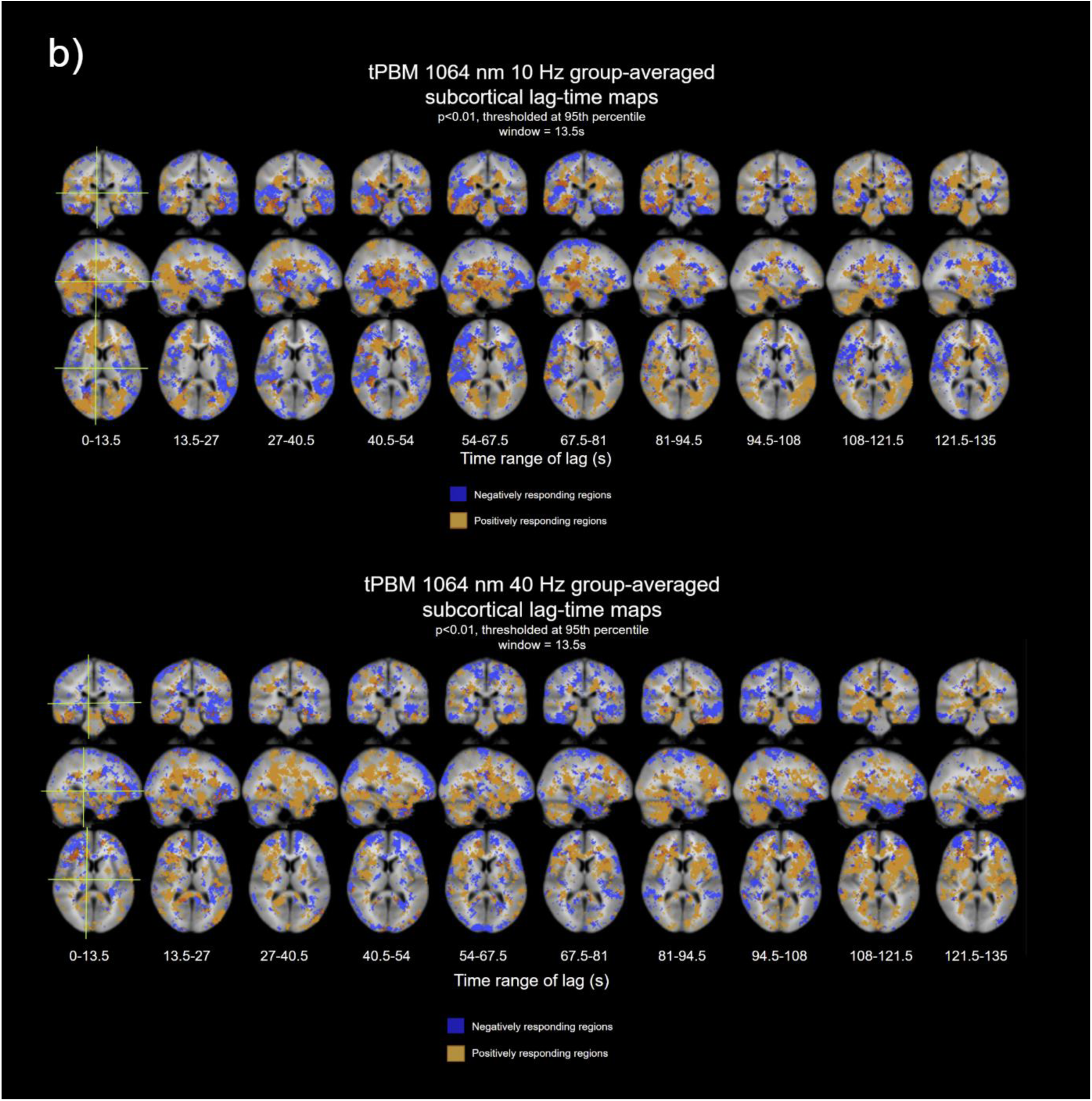
Time-averaged subcortical group-averaged correlation maps for tPBM **a) 808 nm** and **b) 1064 nm** (p<0.01). Maps were thresholded by the 95th percentile of commonality to identify common regions of response at different lag times during the stimulus period. Crosshairs represent coordinates where the sagittal, coronal, and axial slices were taken. Blue and red correspond to regions where significant negative and positive BOLD responses were found (p<0.01).

**Figure 9:**
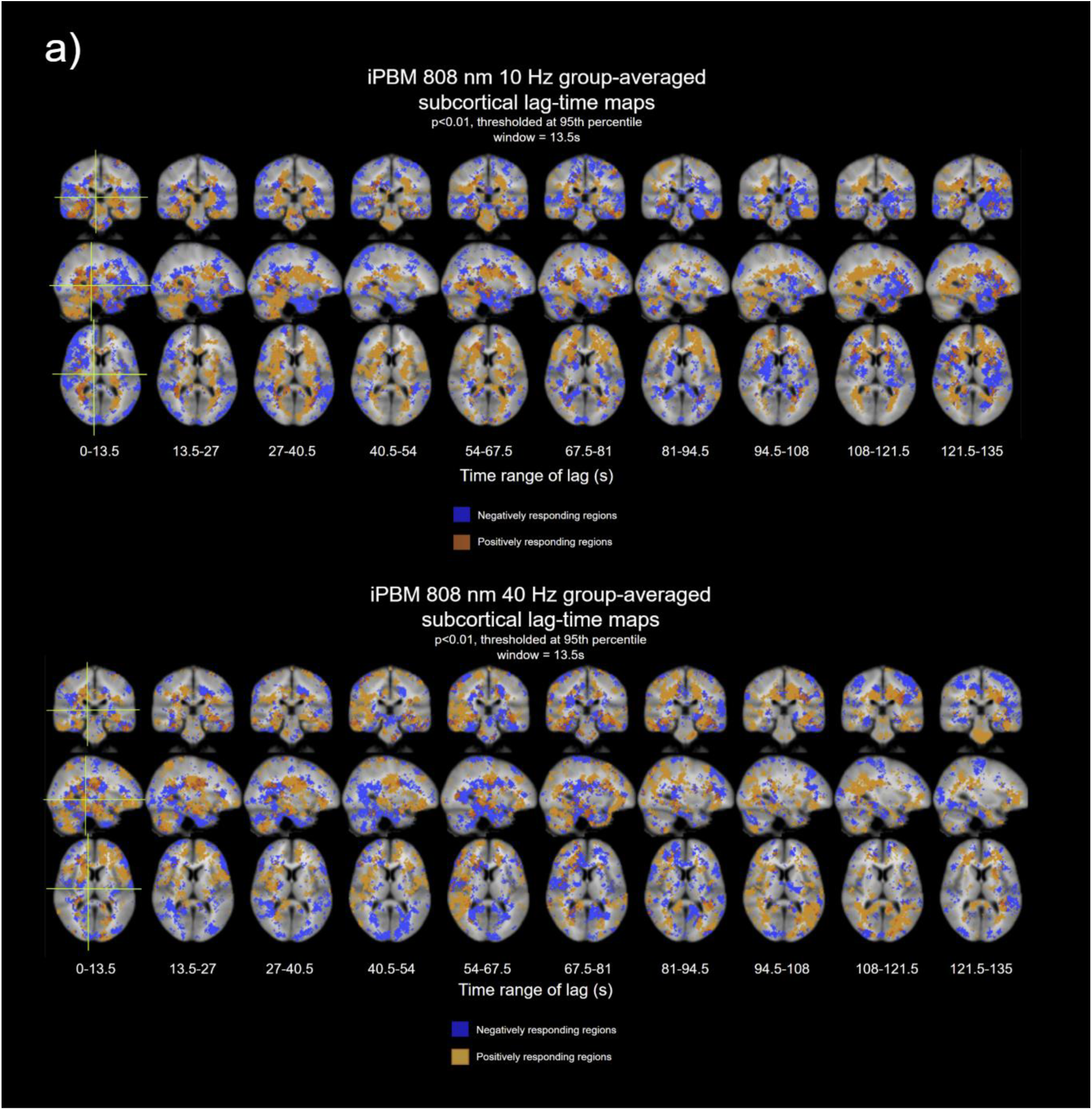
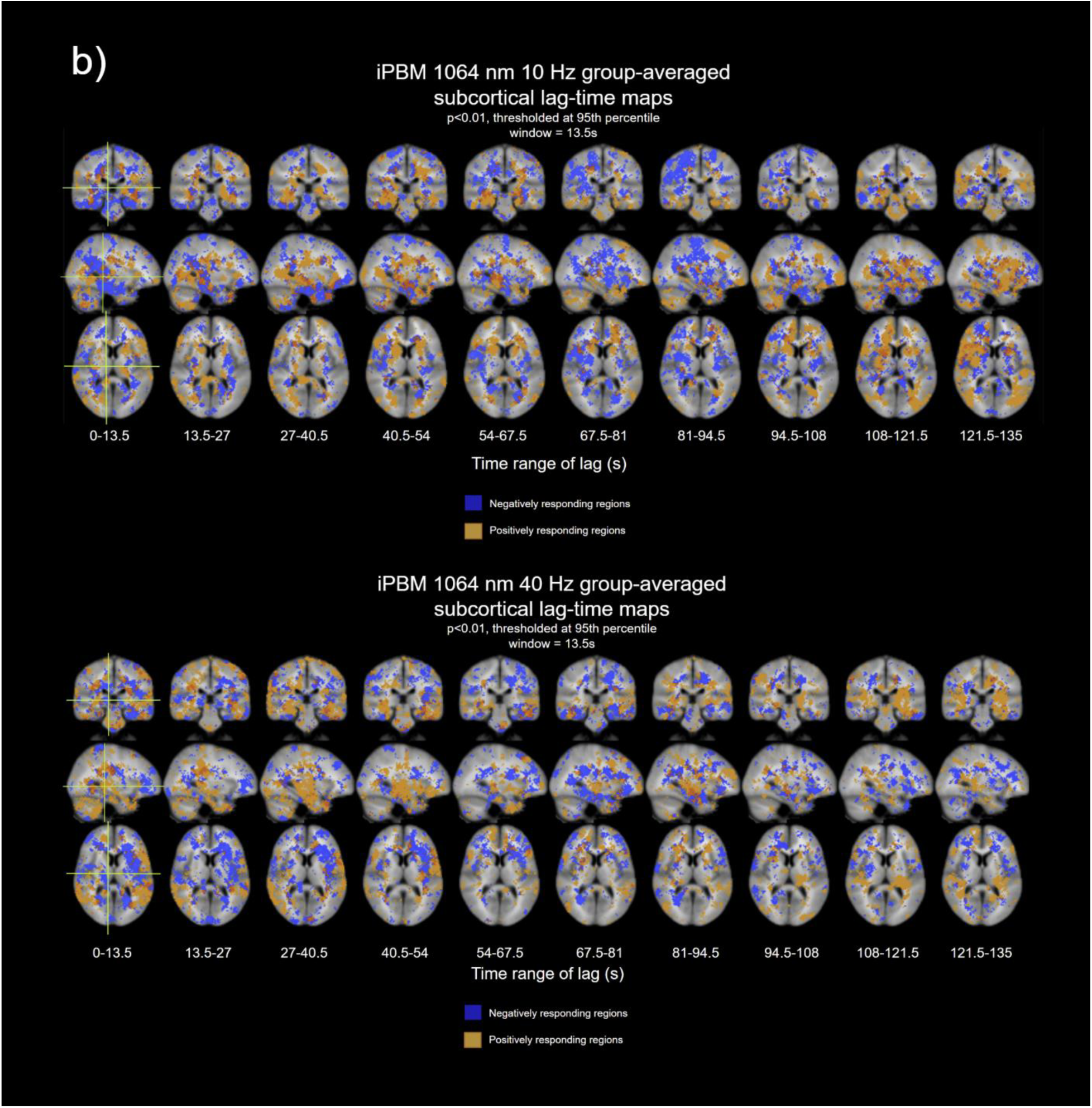
Time-averaged subcortical group-averaged correlation maps for iPBM **a) 808 nm** and **b) 1064 nm** (p<0.01). Maps were thresholded by the 95th percentile of commonality to identify common regions of response at different lag times during the stimulus period. Crosshairs represent coordinates where the sagittal, coronal, and axial slices were taken. Blue and red correspond to regions where significant negative and positive BOLD responses were found (p<0.01).

The volumetric maps show the extent of negative and positive BOLD responses in the subcortical regions in response to both tPBM and iPBM, as well as their global spread across the brain (Figs. 8 and 9, respectively). Generally, both positive and negative BOLD responses are extensive across the white matter.

### Linear mixed effects analysis

**Table 1** summarizes significant online and offline effects revealed by LME analysis.

**Table 3.** Summary of significant parameters revealed from LME analysis. a) ONLINE, during stimulus effects b) OFFLINE, post-stimulus effects. The normalized effect size was calculated by dividing the effect size by the mean of the response variable.

a) Linear mixed effects analysis results on online (stimulus) effects
| Site | Effect | Parameter | Model design | Significance | Normalized Effect Size |
| --- | --- | --- | --- | --- | --- |
| iPBM | Frequency; 40Hz compared to 10Hz | %BOLD online $\tau$, positively responding regions | $R \sim \text{Frequency} + (1\vert \text{Subject})$ | $p < 0.05$ | +0.3027 |

| Site | Effect | Parameter | Model design | Significance | Normalized Effect Size |
| --- | --- | --- | --- | --- | --- |
| tPBM | ITA | %BOLD offline $\tau$, positively responding regions | $R \sim \text{ITA} + (1\vert \text{Subject})$ | $p < 0.05$ | +0.0053 |
| tPBM | ITA | %BOLD offline $\tau$, negatively responding regions | $R \sim \text{ITA} + (1\vert \text{Subject})$ | $p < 0.05$ | +0.0039 |
| iPBM | Frequency; 40Hz compared to 10Hz | %BOLD offline $\tau$, positively responding regions | $R \sim \text{Frequency} + (1\vert \text{Subject})$ | $p < 0.05$ | +0.4213 |

LME analysis found that ITA had significant effects on the dynamics of the offline %BOLD response in tPBM. Higher ITA, and thus lighter skin tone, resulted in a slower return to baseline %BOLD signal in both positively and negatively responding regions. Additionally, the LME analysis also found that 40 Hz pulsation frequency had significant effects in positively responding regions in iPBM. Compared to 10 Hz, 40 Hz resulted in a slower rise in %BOLD signal during stimulus, and a slower return to baseline levels post-stimulus.

## Discussion

This study aimed to characterize, through BOLD-fMRI, the temporal and spatial evolution of the brain response to tPBM and iPBM, highlighting the effects of different light parameters on the dynamics of the BOLD response. To our knowledge, this is the first study of its kind. Our main findings are: (1) tPBM and iPBM both induce a time-varying brain response that corresponds to the timing of the light stimulus, observed as both positive and negative BOLD-fMRI responses in the grey matter; (2) these effects show regional specificity, but are not restricted to the site of stimulation - instead, they are linked to different spatial distributions for tPBM and iPBM delivery; (3) these BOLD effects shift from locations in the anterior brain, proximal to the stimulation site, over to the posterior brain over a period averaging about ∼50 seconds; (4) these effects exhibit both dose and skin tone dependence.

### Interpretation of the positive BOLD-fMRI response

Our study uses BOLD-fMRI to characterize the physiological response to PBM. As the literature has shown that increased production of ATP and NO immediately follow PBM onset (Quirk and Whelan 2020; Barolet et al. 2024), we expected a time-locked positive BOLD response. The observations of PBRs are consistent with past findings using fNIRS in which report that during PBM stimulus, oxidized CCO increases concurrently with concentrations of oxyhemoglobin (Wang et al. 2017, 2016; Tian et al. 2016; Holmes et al. 2019). This is consistent with the assumptions of neurovascular coupling in PBRs, where increased CMRO₂ results in increased inflow of oxygen-rich blood, resulting in a higher BOLD signal. Previous fNIRS studies have shown this clearly in the frontal regions (Wang et al. 2017; Saucedo et al. 2021), with neurovascular coupling enhanced during PBM compared to rest. Therefore, in conjunction with past work, these findings support the current understanding that there is a significant hemodynamic element in the immediate physiological response to PBM, as well as further corroborate evidence that neurovascular coupling is present in driving the neurophysiological response. Nonetheless, although it is generally accepted that positive BOLD activity can represent neural and metabolic activity, the BOLD signal can only act as an indirect marker of neuronal oxygen consumption (Ogawa et al. 1990) due to the complex composition and variables contributing to its signal. The spatiotemporal properties of the BOLD response to stimulus are therefore strongly dependent upon neurovascular coupling.

Unlike these previous studies, we observed the PBR propagating to the whole brain instead of staying in the vicinity of the site of irradiation, a point which will be further discussed later.

### Interpretation of the negative BOLD-fMRI response

Importantly, in addition to PBRs, our results also show persistent negative BOLD responses (NBRs). This is the first time, to our knowledge, that widespread NBRs are observed in response to PBM in conjunction with PBR. Although NBRs are commonly observed in cognitive and clinical studies, it is not as well understood or interpretable as positive BOLD responses (Schridde et al. 2008). BOLD-fMRI is inherently driven by the interplay amongst CBF, cerebral blood volume, and CMRO_2_ (Ogawa et al. 1990), which makes tracing back to any individual substrate difficult. NBR can originate not only due to decreased neural activity, but can also be due to hemodynamic changes that are independent of local neuronal activity, like in the case of blood redistribution, or a mismatch between neural and hemodynamic demands, like in the case of neurovascular decoupling (Moraschi, DiNuzzo, and Giove 2012). Possibly, CMRO_2_ and CBF upregulation in certain regions may be coupled with downregulation in neighbouring regions. It is also possible that an increase in oxidative metabolism in the absence of a blood flow increase produces a NBR ((Schridde et al. 2008; Ogawa et al. 1990).

While the interpretation of the NBR is limited due to the unimodality of our data, certain characteristics of the NBR and its appearance in conjunction with PBR suggest that PBM-induced NBR is caused by inhibition or reduction of local neural activity. An inhibition of neural activity may be caused by astrocyte networks, as recent literature has shown that ATP released from astrocytes may reduce neuronal excitability and signal transmission (Lezmy 2023; Matos et al. 2018; Henriques et al. 2022; Tan et al. 2017). The influence of PBM on astrocytic activity has also been well reported within the literature (Naour et al. 2023; El Massri et al. 2018, 2016; Begum et al. 2013; Blivet et al. 2018; Salehpour et al. 2022; Mitrofanis et al. 2023). Under normal neurovascular coupling, reduced neural activity would lower local CMRO₂, subsequently lowering incoming blood supply, and thereby reducing the %BOLD signal. Tight coupling between PBR and NBR has been observed in past works that compared timings and amplitude of the %BOLD response, suggesting that the coupling mechanism between CCO and the hemodynamic response may be similar (Shmuel et al. 2002, 2006; Liu et al. 2011; Kastrup et al. 2006; Pasley, Inglis, and Freeman 2007). As our results show similar timings and amplitudes for the PBR and NBR (Fig. 8), this makes the theory of NBR originating from neural inhibition plausible.

Although this does not preclude a purely hemodynamic origin of NBR from being possible, our results, at least on a group level, do not demonstrate the level of spatial specificity or coupled activity for adjacent positive and negative regions that would suggest either passive or active mechanisms of blood redistribution during stimulus (Harel et al. 2002; Liu et al. 2011).

### Evolution and extent of the global fMRI response

As mentioned earlier, a novel finding in this work is the propagation of the PBM-fMRI response to distal regions of the brain. A global response in both hemispheres has previously been observed during localized unilateral tPBM irradiation using fNIRS (Tian et al. 2016; Holmes et al. 2019; Wang et al. 2017). Ipsilateral responses in the precuneus from forehead PBM have also been previously noted (Saltmarche et al. 2017; Dmochowski et al. 2020). Several fNIRS studies have reported increased brain connectivity and spectral power of oxyhemoglobin concentration in the medial and lateral prefrontal cortex, the parietal complex, the temporal lobe, and the occipital lobe during and following stimulus applied to the right prefrontal cortex at either 1064 nm or 808 nm wavelength (Truong et al. 2022; Urquhart et al. 2020). Similar increases in brain connectivity in those regions were also reported in studies using EEG and fMRI (Dmochowski et al. 2020; Shahdadian et al. 2022; Naeser et al. 2020; El Khoury, Mitrofanis, and Henderson 2019; Saltmarche et al. 2017; Chao 2019b; Zomorrodi et al. 2019; Urquhart et al. 2020; Argilés et al. 2022). Importantly, while our results are generally aligned with past results, with the reported significantly responding regions remaining consistent, we show how the PBRs and NBRs evolve spatially over the course of the tPBM stimulus.

While differences between protocol groups as well as the effect of stimulation site can be observed, we found recurring patterns in both the locations as well as in the directions of spread of the positive and negative BOLD responses. These patterns of propagation are summarized in Figure 10, in which common regions of response and the general pattern of evolution for all protocols in tPBM and iPBM are summarized. This shift from “early” to “late” responders takes place over ∼90 seconds in most cases, often with remarkable consistency in the specific location and sequence in which the region responds (Figs. 6 & 7). Moreover, while the early iPBM responses are more anterior-inferior, the early tPBM responses appear to be more anterior-superior, as we would expect from simulation studies. Subsequently, however, there is remarkable consistency and conservation between significant nodes in the global propagation of the PBM response across tPBM and iPBM (Fig. 10). Examples of such conserved nodes include the super marginal gyrus, orbitofrontal area, and Broca’s area. Additionally, the direction of the spread in both the medial and lateral regions also remains fairly consistent, with propagation towards the paracentral region and ending at the parieto-occipital regions at later times. The comparability of the iPBM to the tPBM response, despite much lower irradiances used in our iPBM protocol, is consistent with simulation results as well (Cassano et al. 2019; Van Lankveld et al. 2024). The simulations also suggest that in iPBM, as photons do not come into contact with the skull and instead interact with the much thinner and porous cribiform plate, and as there is more air and less cerebrospinal fluid in the intranasal passage than in the forehead path, light could reach the cortex with less attenuation than in the case of tPBM. This may also explain the presence of almost equivalent tPBM and iPBM response amplitudes despite the much lower irradiances used in iPBM.

**Figure 10:**
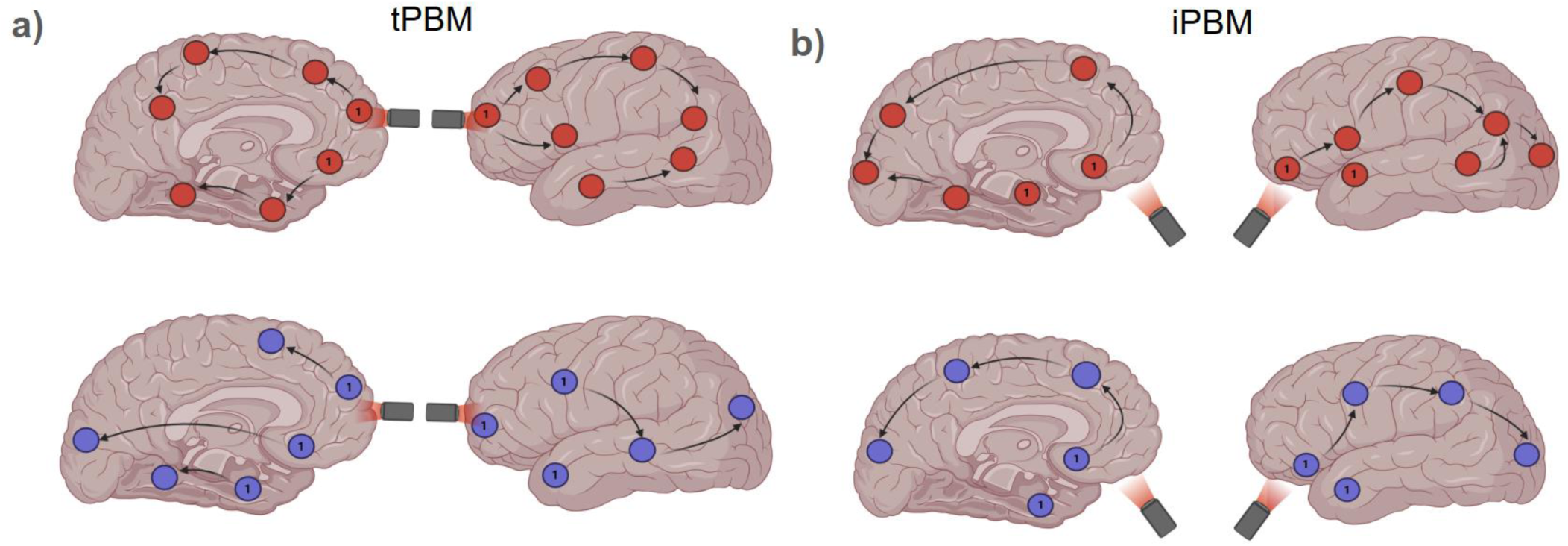
Response evolution patterns in the medial and lateral brain for **(a)** tPBM and **(b)** iPBM. The lasers indicate the sites of irradiation. The red nodes represent regions correlated with a positive BOLD response, while the blue nodes represent regions correlated with a negative BOLD response. Nodes marked with 1 represent regions found to be initial responders to the stimulus. Arrows indicate the direction of spread and connections between other nodes, as indicated by ordering the lag times.

The spatial selectivity of the response and consistency of the diffuse pattern of response suggest a common underlying mechanism driving the global PBM response. The involvement of brain networks could be a potential source of this response. Past work has identified several functional networks, such as the default-mode network, the fronto-parietal network, and the dorsal attention network, to undergo alterations and see increases in functional connectivity during and after PBM treatment (Naeser et al. 2020; El Khoury, Mitrofanis, and Henderson 2019; Saltmarche et al. 2017; Chao 2019a; Zomorrodi et al. 2019; Urquhart et al. 2020; Argilés et al. 2022). Ours is the first study to demonstrate temporal evolution in terms of potential network involvement. In our results, we can see nodal regions in each network respond to PBM at different lag times. For example, in the 1064 nm 10 Hz tPBM protocol group, we can see that the PBR begins at the medial prefrontal cortex at the start of stimulus (see Fig 6b). Gradually, we see the PBR spreading to the precuneus and hippocampus at 54-67.5 s and ∼104 s lag times, respectively, which correlates with regions within the default-mode network. PBR at the medial prefrontal cortex and the precuneus can be observed in almost all protocols in tPBM and iPBM. Similar lagged response patterns related to the fronto-parietal network can also be seen in 808 nm 40 Hz protocol groupings in tPBM and iPBM as well, with response beginning at the rostral lateral and dorsolateral prefrontal cortex, and later spreading to the anterior inferior parietal lobule and posterior inferior temporal lobe.

The mode of response propagation is akin to findings in other neuromodulation studies, such as those investigating the dynamics of transcranial magnetic stimulation (Valero-Cabre, Payne, and Pascual-Leone 2007; Valero-Cabré et al. 2005; Siebner et al. 2009; Ruff, Driver, and Bestmann 2009; Ferreri et al. 2011; Lisanby and Belmaker 2000; Fox et al. 2012). In those works, the spread of activation has been suggested to be facilitated by intra and inter-hemispheric cortico-cortical connectivity between regions (Lee et al. 2003; Komssi and Kähkönen 2006). However, the speed of spread across networks in our results appears slower than would be warranted by a purely neuronal mechanism. Thus, we also considered a vascular mechanism unrelated to neurovascular coupling. Recent research has shown that PBM induces significant changes to endogenic oscillations (0.003–0.02 Hz) during stimulus (Truong et al. 2022), which have been shown to be correlated with increased vascular endothelial metabolic activity, such as the synthesis and release of vasodilators like NO and endothelial-derived hyperpolarizing factors, which would increase local cerebral blood flow and increase the strength of the BOLD signal (Rubanyi 1991). Such infra-slow vascular fluctuations have been reported to propagate across the brain in specific network-dependent patterns in both cortical and subcortical structures, which is consistent with our results (Raut et al. 2021; Bright et al. 2020; Nanayakkara et al. 2025). Additionally, the propagation within and across networks has been shown to possess lags between regions of around ∼10 s to ∼40 s, which would be slow enough to be consistent with the lag times we observed. Active propagation through hyperpolarizing currents and ion pulses at the gap junctions of endothelial cells may also be involved (Broggini et al. 2024). As such, a potential vascular mechanism of driving the spread of global BOLD response from the initial stimulation site could be that 1) photo-oxidation of CCO in sites proximal to the stimulation site enhances local CCO redox metabolism and ATP synthesis, and increases in the bioavailability of NO; 2) this results in changes to local vascular endothelial activity, smooth muscle relaxation, and increases in cerebral blood flow; 3) these vascular changes are reflected in infra-slow oscillations in the vascular network that proceed to propagate throughout the rest of the brain in a network-dependent fashion through existing vascular networks; 4) the subsequently induced vascular endothelial metabolic activity in distant regions result in the synthesis and release of vasodilating factors in these distal reaches of the vascular networks.

An additional point of interest is that our results also show extensive and dynamic PBR and NBR in the subcortical regions, with their strength and location evolving across different time windows (Fig. 8 & **9**). A potential neuronal mechanism, as suggested in recent literature, is that calcium-dependent ATP release by white matter astrocytes regulates and controls the conduction velocity of axonal action potentials (Hamilton et al. 2008; Lezmy et al. 2021). Normally triggered by a rise in intracellular calcium levels, ATP release from fibrous astrocytes leads to the binding of extracellular adenosine to neuronal receptors, triggering hyperpolarization in potassium/sodium hyperpolarization-activated cyclic nucleotide-gated ion channel 2 (HCN2), which ultimately dampens the speed of impulse propagation across networks (Lezmy et al. 2021). Theorized to serve as a potential brake to long-range neural activity, this could explain the extensive distribution of PBR and NBR in the white matter, as well as demonstrate how neuronal networks might play a role in the lagged and slowed propagation of the PBM response across the brain. However, BOLD-fMRI in the white matter is still controversial, and it remains unclear whether vascular mechanisms independent of neuronal activity contributed to these dynamics.

### Dose dependence of fMRI response dynamics

The likely mechanisms of the PBM-fMRI responses can also be gleaned from their dose-dependency. Our LME results reveal a dependence of the PBM response dynamics on laser pulsation frequency. We observed a frequency dependence in the PBR in iPBM (both online and offline). Interestingly, compared to 10 Hz, stimulation at 40 Hz frequency resulted in a slower rise of the BOLD signal upon stimulus onset, as well as a slower return to baseline levels in the post-stimulus period. This frequency-dependent speed of the transient BOLD response may be determined by the interplay between metabolic and hemodynamic transients (Chen and Pike 2009). For instance, a more sluggish BOLD post-stimulus response could be due to either a sustained elevation in blood flow or an early reduction in oxidative metabolism. Although there is sparse literature on the 10 Hz pulsation in forehead tPBM, prior research has shown that tPBM at 40 Hz increased alpha, beta, and gamma power while reducing power in the delta and theta frequency bands (Zomorrodi et al. 2019). tPBM at 40 Hz has also been reported to alter EEG microstates and complexity (Zhang et al. 2021). As different EEG frequency bands have been shown to exhibit distinct coupling with hemodynamic activity in neurovascular coupling studies that varied by region of interest, task and disease state, it is possible that different regions in the brain respond to PBM depending on the initial stimulation site, and modulation of the alpha and gamma powers in that region led to changes in neurovascular coupling, leading to differences in the offline BOLD response (*New Horizons in Neurovascular Coupling: A Bridge between Brain Circulation and Neural Plasticity: Volume 225* 2016; Lecrux and Hamel 2016; Girouard and Iadecola 2006; Pinti et al. 2021; Chiarelli et al. 2021; Chalak et al. 2017).

Additionally, our LME results revealed that individual skin tones affected the offline characteristics of both the PBR and NBR in tPBM. We found that higher ITA, and therefore lighter skin tone, resulted in a slower BOLD return to baseline. It is possible that higher energy deposition, in the case of lighter skin tones, could result in higher levels of NO as well as higher rates of ATP production and oxygen consumption by CCO, co-creating a more gradual PBR rise. It should be noted that skin tone could also be coincidentally correlated with other subject-specific variations, such as the distance from the laser to the brain, but in our LME analysis, no significant correlation was found for this effect.

### Correlation-based versus non-correlation-based analysis

While our results are derived from correlation-based analysis, which is a more novel and uncommon analytical method in fMRI, our results do align and are comparable with other work that does not use correlation-based methods. For example, our lab has previously used independent component analysis (ICA) to analyze the BOLD-fMRI and CBF response elicited by PBM and has found that the response shows regional stimulation parameter dependence with regions closer to the site of stimulation responding differentially to wavelength and frequency (Van Lankveld et al. 2024). Specifically, they reported shorter wavelengths (808 nm) and higher frequencies (40 Hz) promoting an increase in BOLD and CBF signals, compared to deeper structures such as the amygdala-hippocampus, which respond to longer wavelengths and intermediate irradiances. Along with the results derived from our approach in this study, a combination of correlation and non-correlation analyses could yield additional interesting insights towards the regional dependence and evolution of the neurophysiological response.

### Limitations and future work

One caveat towards the results of this work is the subject pool size and the large range of intersubject differences and variability observed in the BOLD response. Increasing the subject pool size would improve the replicability and generalizability of the results. Another caveat of this work to consider is that this study only considers BOLD-fMRI and lacks more quantitative and direct measures of the brain’s physiological state, like CBF. Finally, this study lacks any form of MRI-independent measures to assess the meaning of our finding of the negative BOLD response. Because the BOLD signal is an indirect marker of brain metabolic activity, there could be many factors and interpretations that could result in a negative BOLD signal. Whether or not the negative BOLD response arises due to vascular or neural factors is a question that cannot be answered only through fMRI. Therefore, to gain a more accurate understanding of what is occurring in regions that demonstrate a negative BOLD response, additional measures would be insightful.

### Conclusion

In this study, we investigated the effects of near-infrared light and the impact of stimulation site on the dynamics of the neurophysiological response to PBM. Our results illustrate how the evolution of the global BOLD response during PBM treatment is influenced not only by the stimulation site but also by skin tone and laser parameters. Additionally, we observe co-localizations between positive and negative BOLD-fMRI responses in both the grey and white matter, which suggest that functional networks facilitate the propagation of the physiological response to PBM in the brain, irrespective of stimulation site. To our knowledge, this is the first study investigating the dose-dependent dynamics of the PBM response using BOLD-fMRI.

## Acknowledgements

This work was supported by funding from the Ontario Centre for Innovation, the Natural Sciences and Engineering Research Council (NSERC) of Canada and Vielight Inc.

## Statement of conflict

Vielight Inc. supplied the laser equipment and partial funding, as part of the NSERC Alliance Program and the Ontario Centre for Innovation Program. Vielight Inc. was not involved in author equity, consultancy, or patent status by the authors, study design, analysis or manuscript preparation.

## Notes

### Competing Interest Statement

The authors have declared no competing interest.

