## Supplementary Materials for "Real-time spatial evolution of the fMRI response to photobiomodulation in the healthy human brain"

### 1. Skin-tone measurements and Individual Typological Angle (ITA)

Skin-tone was quantified using a CM-600D spectrophotometer (Konica Minolta, Tokyo, Japan). The device outputs colorimetric values in the CIE $L^*$ $b^*$ color space, where $L^*$ represents luminance (0 = black, 100 = white), and $b^*$ represents the blue-yellow axis. These $L^*$ and $b^*$ values were used to compute individual topography angles (ITA), a skin pigmentation index resulting from both eumelanin and pheomelanin, and it is calculated with the following formula:

$$\textbf{\textit{ITA = [arctan(L*-50)/b*)]*180/}}\boldsymbol{\pi} \qquad (S1)$$

Using a reference of L = 50, the ITA is the angle formed by the $L^*$ (luminance) and $b^*$ (blue-yellow) axes, with higher ITA values corresponding to lighter skin tones. For each participant, six 8 mm diameter measurement samples were taken of the right forehead of each subject prior to PBM laser placement, and the average was taken as the individual's ITA measurement. Zero and white calibration were performed before each use to ensure accuracy of the calculations.

Using these ITA measurements, participants can be grouped into three skin-tone groups based on custom ITA thresholds (**Figure S1a**, **Table S1**), consistent with previous literature (Jung et al. 2023). We also ensured an even ratio of males and females within each skin-tone group (Groups 1, 2, 3). Continuous ITA values were retained for all statistical analyses to better capture individual-level variation in skin light absorption during PBM. **Figure S1b** depicts the distribution of ITA values of the participants within our study.

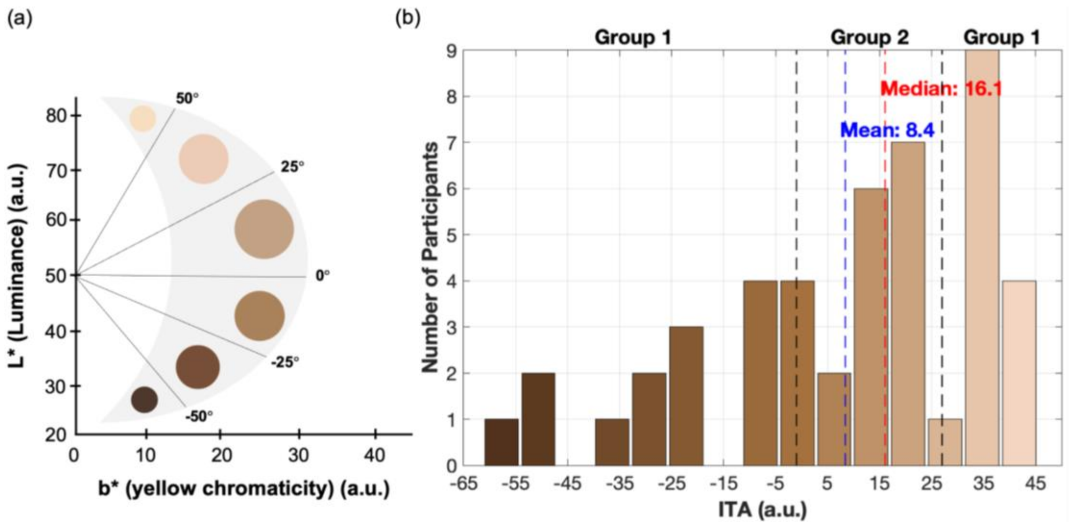


**Figure S1: ITA quantification and distribution. (a)** ITA as a quantification of skin tone. **(b)** Participant distribution across ITAs (median ± range = 16.1 ± 106.7), with the mean and median in blue and red respectively, and group (Groups 1, 2, 3) boundaries in black.

**Table S1: Custom ITA thresholds for different skin-tone groups used in this work.**

| Skin Tone Group | ITA Range |
|---|---|
| Group 1 | 27° < ITA ≤ 55° |
| Group 2 | -1° < ITA ≤ 27° |
| Group 3 | ITA ≤ -1° |

### 2. MR Thermometry

We utilized an MR thermometry sequence that exploits the temperature sensitivity of the resonance frequency in order to scan and collect data on whether or not there were significant thermal effects during and post-PBM stimulus. MR thermometry data was collected in the highest irradiance subgroup at 1064 nm across 30 subjects.

The MR thermometry sequence followed the same stimulus paradigm as the BOLD-fMRI sequence shown in **Figure 2**. The temperature was measured in the illuminated region shown in **Figure S2**. The MRI phase and magnitude images were processed to estimate temperature changes induced by PBM using the Proton Resonance Frequency (PRF) shift method. A custom preprocessing script was used for brain extraction, motion correction, drift correction, artifact removal (4 standard deviations above the average), phase radian conversion, unwrapping corrected for phase aliasing to produce a continuous phase map. Temperature changes (ΔT) were then calculated as follows, with all symbols defined in **Table S2**.

$$\Delta T = \frac{\Delta\phi}{\alpha\gamma\beta_0 TE} \qquad \text{(S2)}$$

Temperature changes were calculated on a frame-by-frame basis, using Equation S2 and the following parameters shown in **Table S2**.

**Table S2: Summary of MR thermometry parameters and used in temperature change calculations.**

| Symbol | Parameter | Values | Units |
|---|---|---|---|
| ΔT | Frame by frame temperature change | | °C |
| Δ | Phase change from baseline | 0 : 4095 | radians |
| γ | Gyromagnetic ratio of hydrogen | $2\pi \cdot 42.58 x 10^6$ | rad/s/T |
| α | PRF thermal coefficient | $0.01 x 10^{-6}$ | ppm/°C |
| $\beta_0$ | Main magnetic field strength | 3 | Tesla |
| TE | Echo time | 0.007398s | seconds |

Temperature time courses were extracted from identified regions of interest and were analyzed separately for the two frequency groups: 10 Hz (N=16) and 40 Hz (N=14) (see **Figure S2, S3**). The temperature of each subject was recorded per frame, and then the mean temperature change was calculated to assess the group-level response. The standard error was also calculated to provide an estimate of variability between subjects. A paired sample t-test was performed to compare the mean temperature during the pre-stimulus recording (minutes 0-4) and PBM stimulus (minutes 4-8), to determine if the mean temperature during PBM was significantly different than the mean temperature during the pre-stimulus recording period.

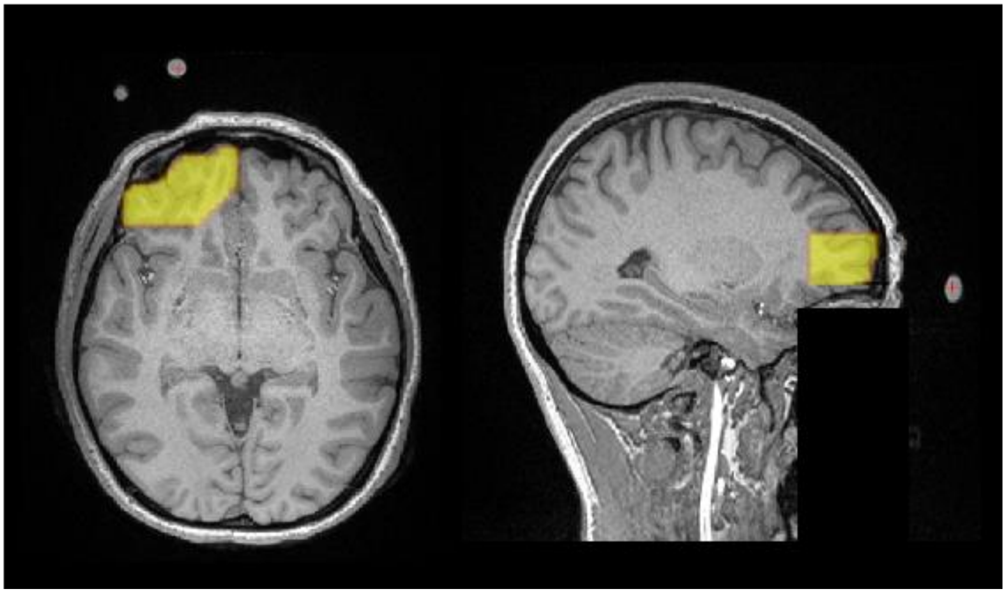

**Figure S2: MR Thermometry region of interest.** The vitamin E capsules were used as fiducial markers to indicate the laser location. Neural tissue in closest proximity to these markers was manually segmented for each participant, and this region was used for MR thermometry analysis.

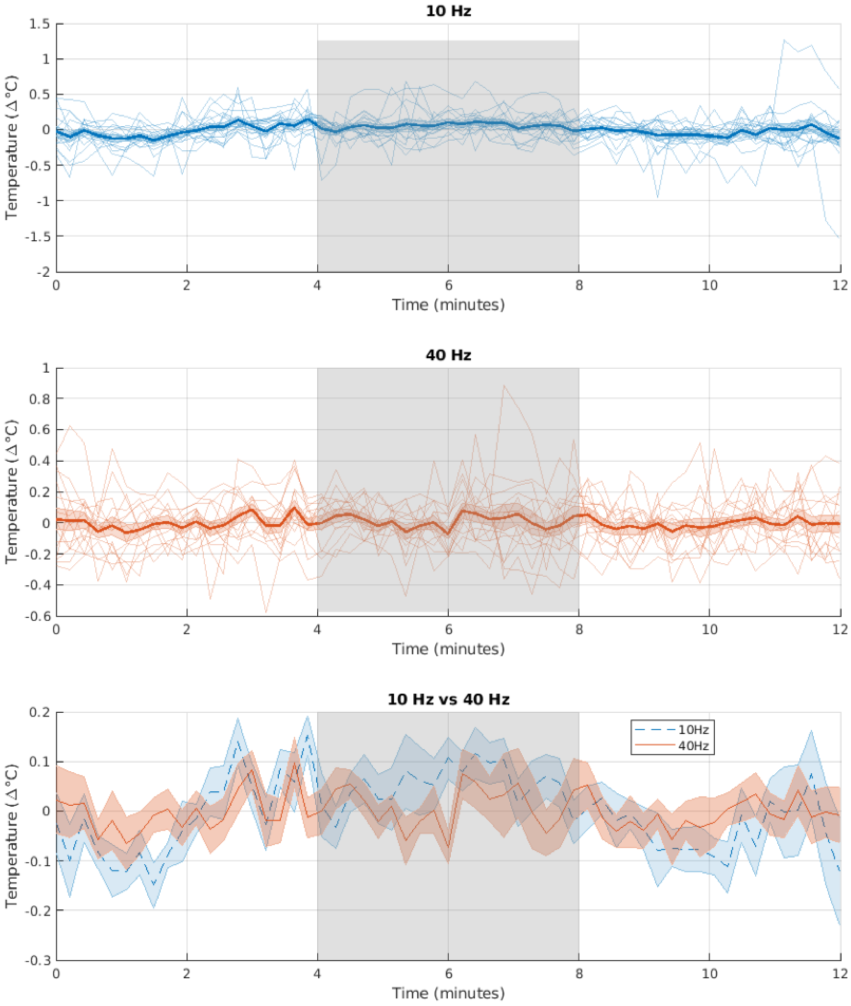


**Figure S3. Temperature changes** (Δ°C) **over time during tPBM at 10 Hz** (top), **40 Hz** (middle), **and their comparison** (bottom). Shaded regions indicate the stimulation window (4–8 minutes). Thin lines represent individual subject responses; bold lines show group averages with standard error shading. The comparison plot emphasizes mean temperature trends for both frequencies, revealing no significant differences in temperature during the stimulation periods.

Group-level statistical comparison showed that the average temperature change was minimal and remained stable over the entire 12 minutes ($p = 0.13$ and 0.76, comparing PBM to pre-PBM for 10 Hz and 40 Hz stimulation, respectively).

### 3. Linear Mixed Effects Analysis: Results

**Table S3:** Summary of all results of the linear mixed effects analysis, organized as a) tPBM online effects; b) tPBM offline effects; c) iPBM online effects; d) iPBM offline effects, listing the coefficient estimate for the effect (Estimate), standard error (SE), the associated t-statistic value (tStat), the degrees of freedom (DOF), the p-value (pValue), the FDR-corrected p-value (FDR_pValue), and the lower and upper bounds of the confidence interval (CI_Lower and CI_Upper).

### a) tPBM Linear Mixed Effects analysis results on online (stimulus) effects

| Effect | Parameter | fitlme Formula | Estimate | SE | tStat | DOF | pValue | FDR_pValue | CI_Lower | CI_Upper |
|---|---|---|---|---|---|---|---|---|---|---|
| ITA | %BOLD online τ, positive response | Parameter~ITA+(1\|Subject) | 0.089791 | 0.014017 | 6.405864 | 152 | 0.210453 | 0.276486 | 0.062335 | 0.117250 |
| ITA | %BOLD online τ, negative response | Parameter~ITA+(1\|Subject) | 0.221767 | 0.059206 | 3.745684 | 144 | 0.295233 | 0.350338 | 0.105716 | 0.337804 |
| ITA | 3-minute mean, positive response | Parameter~ITA+(1\|Subject) | -0.061599 | 0.011216 | -5.492065 | 176 | 0.067211 | 0.112822 | -0.083573 | -0.039607 |
| ITA | 3-minute mean, negative response | Parameter~ITA+(1\|Subject) | -0.005764 | 0.011409 | -0.505215 | 175 | 0.087572 | 0.158369 | -0.028122 | 0.016602 |
| Wavelength; 1064nm compared to 808nm | %BOLD online τ, positive response | Parameter~Wavelength+(1\|Subject) | 0.166173 | 0.018980 | 8.755163 | 152 | 0.291477 | 0.340537 | 0.128969 | 0.203371 |
| Wavelength; 1064nm compared to 808nm | %BOLD online τ, negative response | Parameter~Wavelength+(1\|Subject) | -0.045960 | 0.046553 | -0.987262 | 144 | 0.096643 | 0.157253 | -0.137204 | 0.045284 |
| Wavelength; 1064nm compared to 808nm | 3-minute mean, positive response | Parameter~Wavelength+(1\|Subject) | 0.135624 | 0.029181 | 4.647681 | 176 | 0.255753 | 0.311961 | 0.078427 | 0.192813 |
| Wavelength; 1064nm compared to 808nm | 3-minute mean, negative response | Parameter~Wavelength+(1\|Subject) | 0.132397 | 0.090845 | 1.457394 | 175 | 0.216954 | 0.284904 | -0.045578 | 0.310358 |
| Frequency; 40Hz compared to 10Hz | %BOLD online τ, positive response | Parameter~Frequency+(1\|Subject) | 0.032026 | 0.031327 | 1.022313 | 152 | 0.060442 | 0.112479 | -0.029381 | 0.093421 |
| Frequency; 40Hz compared to 10Hz | %BOLD online τ, negative response | Parameter~Frequency+(1\|Subject) | -0.109764 | 0.033735 | -3.253713 | 144 | 0.098891 | 0.160147 | -0.175881 | -0.043639 |
| Frequency; 40Hz compared to 10Hz | 3-minute mean, positive response | Parameter~Frequency+(1\|Subject) | 0.374937 | 0.094635 | 3.961927 | 176 | 0.443693 | 0.490783 | 0.189445 | 0.560415 |
| Frequency; 40Hz compared to 10Hz | 3-minute mean, negative response | Parameter~Frequency+(1\|Subject) | 0.029090 | 0.014804 | 1.965009 | 175 | 0.073453 | 0.142542 | 0.000074 | 0.058106 |
| Power; 150mw/cm2 compared to 100mw/cm2 | %BOLD online τ, positive response | Parameter~Power+(1\|Subject) | 0.005473 | 0.051448 | 0.106379 | 152 | 0.061672 | 0.119256 | -0.095368 | 0.106308 |
| Power; 150mw/cm2 compared to 100mw/cm2 | %BOLD online τ, negative response | Parameter~Power+(1\|Subject) | 0.019015 | 0.008146 | 2.334274 | 144 | 0.072647 | 0.127978 | 0.003044 | 0.034976 |
| Power; 150mw/cm2 compared to | 3-minute mean, positive response | Parameter~Power+(1\|Subject) | 0.092113 | 0.028997 | 3.176639 | 176 | 0.210574 | 0.276426 | 0.035276 | 0.148944 |

| | | | | | | | | | | |
|---|---|---|---|---|---|---|---|---|---|---|
| 100mw/cm2 | | | | | | | | | | |
| Power; 150mw/cm2 compared to 100mw/cm2 | 3-minute mean, negative response | Parameter~Power+(1\|Subject) | -0.145537 | 0.058882 | -2.47167 | 175 | 0.095538 | 0.146002 | -0.030121 | -0.260939 |
| Power; 200mw/cm2 compared to 200mw/cm2 | %BOLD online τ, positive response | Parameter~Power+(1\|Subject) | -0.026053 | 0.025029 | -1.04091 | 152 | 0.127278 | 0.108335 | 0.023007 | -0.075107 |
| Power; 200mw/cm2 compared to 200mw/cm2 | %BOLD online τ, negative response | Parameter~Power+(1\|Subject) | -0.111125 | 0.021481 | -5.17318 | 144 | 0.399588 | 0.468891 | -0.069017 | -0.153223 |
| Power; 200mw/cm2 compared to 100mw/cm2 | 3-minute mean, positive response | Parameter~Power+(1\|Subject) | -0.036141 | 0.010201 | -3.54289 | 176 | 0.136439 | 0.190543 | -0.016146 | -0.056134 |
| Power; 200mw/cm2 compared to 100mw/cm2 | 3-minute mean, negative response | Parameter~Power+(1\|Subject) | -0.096944 | 0.021396 | -4.53094 | 175 | 0.844418 | 0.903398 | -0.055004 | -0.138876 |
| Nasal distance (mm) | %BOLD online τ, positive response | Parameter~Distance+(1\|Subject) | 0.279956 | 0.021061 | 13.29263 | 152 | 0.128897 | 0.194585 | 0.321823 | 0.238627 |
| Nasal distance (mm) | %BOLD online τ, negative response | Parameter~Distance+(1\|Subject) | 0.161424 | 0.032277 | 5.001208 | 144 | 0.140565 | 0.192281 | 0.224683 | 0.098157 |
| Nasal distance (mm) | 3-minute mean, positive response | Parameter~Distance+(1\|Subject) | -0.93484 | 0.073652 | -12.6927 | 176 | 0.191344 | 0.25323 | -0.790482 | -1.079198 |
| Nasal distance (mm) | 3-minute mean, negative response | Parameter~Distance+(1\|Subject) | 0.012340 | 0.053778 | 0.229462 | 175 | 0.264884 | 0.320460 | 0.117745 | -0.093065 |

### b) tPBM Linear Mixed Effects analysis results on offline (stimulus) effects

| Effect | Parameter | fitlme Formula | Estimate | SE | tStat | DOF | pValue | FDR_pValue | CI_Lower | CI_Upper |
|---|---|---|---|---|---|---|---|---|---|---|
| ITA | %BOLD offline τ, positive response | Parameter~ITA+(1\|Subject) | 0.137101 | 0.054440 | 2.518387 | 152 | 0.000764 | 0.013150 | 0.029229 | 0.244770 |
| ITA | %BOLD offline τ, negative response | Parameter~ITA+(1\|Subject) | 0.110342 | 0.038656 | 2.85446 | 146 | 0.000126 | 0.005070 | 0.033811 | 0.0050701 |
| ITA | 3-minute mean, positive response | Parameter~ITA+(1\|Subject) | 0.004327 | 0.000656 | 6.596037 | 175 | 0.061756 | 0.300112 | 0.003014 | 0.005586 |
| ITA | 3-minute mean, negative response | Parameter~ITA+(1\|Subject) | 0.008100 | 0.000299 | 27.0903 | 169 | 0.087499 | 0.388066 | 0.007513 | 0.008687 |
| Wavelength; 1064nm compared to 808nm | %BOLD offline τ, positive response | Parameter~Wavelength+(1\|Subject) | 0.237441 | 0.048323 | 4.913623 | 152 | 0.340025 | 0.395667 | 0.142693 | 0.332107 |
| Wavelength; 1064nm compared to 808nm | %BOLD offline τ, negative response | Parameter~Wavelength+(1\|Subject) | -0.021465 | 0.006883 | -3.11855 | 146 | 0.053183 | 0.131932 | -0.034892 | -0.007911 |
| Wavelength; 1064nm compared to 808nm | 3-minute mean, positive response | Parameter~Wavelength+(1\|Subject) | 0.192482 | 0.060292 | 3.192497 | 175 | 0.315590 | 0.621608 | 0.074232 | 0.310568 |
| Wavelength; 1064nm compared to 808nm | 3-minute mean, negative response | Parameter~Wavelength+(1\|Subject) | 0.141336 | 0.041827 | 3.379061 | 169 | 0.258726 | 0.306584 | 0.059333 | 0.223267 |
| Frequency; 40Hz compared to 10Hz | %BOLD offline τ, positive response | Parameter~Frequency+(1\|Subject) | 0.002194 | 0.000453 | 4.843267 | 152 | 0.060653 | 0.344684 | 0.001213 | 0.002987 |
| Frequency; 40Hz compared to 10Hz | %BOLD offline τ, negative response | Parameter~Frequency+(1\|Subject) | -0.031272 | 0.008045 | -3.88713 | 146 | 0.046145 | 0.278993 | -0.046978 | -0.015438 |
| Frequency; 40Hz compared to 10Hz | 3-minute mean, positive response | Parameter~Frequency+(1\|Subject) | 0.390646 | 0.031185 | 12.52673 | 175 | 0.474485 | 0.529072 | 0.329477 | 0.451723 |
| Frequency; 40Hz compared to 10Hz | 3-minute mean, negative response | Parameter~Frequency+(1\|Subject) | 0.0783224 | 0.047334 | 1.654675 | 169 | 0.222462 | 0.263931 | -0.014452 | 0.171095 |

| | | | | | | | | | | |
|---|---|---|---|---|---|---|---|---|---|---|
| Power; 150mw/cm2 compared to 100mw/cm2 | %BOLD offline τ, positive response | Parameter~Power+(1\|Subject) | 0.0067402 | 0.004338 | 1.553757 | 152 | 0.387145 | 0.414137 | -0.001804 | 0.015203 |
| Power; 150mw/cm2 compared to 100mw/cm2 | %BOLD offline τ, negative response | Parameter~Power+(1\|Subject) | 0.0813501 | 0.008926 | 9.113836 | 146 | 0.117993 | 0.171188 | 0.063855 | 0.098845 |
| Power; 150mw/cm2 compared to 100mw/cm2 | 3-minute mean, positive response | Parameter~Power+(1\|Subject) | 0.124002 | 0.033437 | 3.708526 | 175 | 0.166336 | 0.230563 | 0.054463 | 0.185537 |
| Power; 150mw/cm2 compared to 100mw/cm2 | 3-minute mean, negative response | Parameter~Power+(1\|Subject) | -0.094334 | 0.057772 | -1.63287 | 169 | 0.346191 | 0.396223 | -0.207530 | 0.018933 |
| Power; 200mw/cm2 compared to 200mw/cm2 | %BOLD offline τ, positive response | Parameter~Power+(1\|Subject) | 0.012456 | 0.004301 | 2.896071 | 152 | 0.164347 | 0.220072 | 0.003971 | 0.020830 |
| Power; 200mw/cm2 compared to 200mw/cm2 | %BOLD offline τ, negative response | Parameter~Power+(1\|Subject) | -0.092242 | 0.008841 | -10.4334 | 146 | 0.318272 | 0.388318 | -0.109530 | -0.074873 |
| Power; 200mw/cm2 compared to 100mw/cm2 | 3-minute mean, positive response | Parameter~Power+(1\|Subject) | 0.03627 | 0.008311 | 4.364096 | 175 | 0.292526 | 0.346145 | 0.019913 | 0.052494 |
| Power; 200mw/cm2 compared to 100mw/cm2 | 3-minute mean, negative response | Parameter~Power+(1\|Subject) | -0.092383 | 0.006726 | -13.7352 | 169 | 0.118335 | 0.176893 | -0.105568 | -0.079203 |
| Nasal distance (mm) | %BOLD offline τ, positive response | Parameter~Distance+(1\|Subject) | 0.325646 | 0.022191 | 14.67469 | 152 | 0.167436 | 0.233282 | 0.282106 | 0.369094 |
| Nasal distance (mm) | %BOLD offline τ, negative response | Parameter~Distance+(1\|Subject) | -0.094362 | 0.030947 | -3.04915 | 146 | 0.496343 | 0.547668 | -0.154962 | -0.033641 |
| Nasal distance (mm) | 3-minute mean, positive response | Parameter~Distance+(1\|Subject) | -0.913552 | 0.068212 | -13.3928 | 175 | 0.150284 | 0.212035 | -1.047211 | -0.779833 |
| Nasal distance (mm) | 3-minute mean, negative response | Parameter~Distance+(1\|Subject) | 0.066974 | 0.049218 | 1.360762 | 169 | 0.125753 | 0.181327 | -0.029571 | 0.163367 |

### c) iPBM Linear Mixed Effects analysis results on online (stimulus) effects

| Effect | Parameter | fitlme Formula | Estimate | SE | tStat | DOF | pValue | FDR_pValue | CI_Lower | CI_Upper |
|---|---|---|---|---|---|---|---|---|---|---|
| ITA | %BOLD online τ, positive response | Parameter~ITA+(1\|Subject) | 2.959792 | 1.325858 | 2.23236 | 148 | 0.133913 | 0.200954 | 0.361110 | 5.558474 |
| ITA | %BOLD online τ, negative response | Parameter~ITA+(1\|Subject) | -1.589349 | 2.790979 | -0.56946 | 147 | 0.385756 | 0.624056 | -7.059668 | 3.880970 |
| ITA | 3-minute mean, positive response | Parameter~ITA+(1\|Subject) | 0.140454 | 0.094874 | 1.480427 | 171 | 0.151632 | 0.452163 | -0.045495 | 0.326395 |
| ITA | 3-minute mean, negative response | Parameter~ITA+(1\|Subject) | -0.269334 | 0.018106 | -14.8754 | 175 | 0.189585 | 0.245164 | -0.304818 | -0.233842 |
| Wavelength; 1064nm compared to 808nm | %BOLD online τ, positive response | Parameter~Wavelength+(1\|Subject) | 5.679464 | 2.579016 | 2.202183 | 148 | 0.476751 | 0.555449 | 0.624593 | 10.734335 |
| Wavelength; 1064nm compared to 808nm | %BOLD online τ, negative response | Parameter~Wavelength+(1\|Subject) | 6.002384 | 2.828953 | 2.121769 | 147 | 0.196552 | 0.502518 | 0.457636 | 11.547132 |
| Wavelength; 1064nm compared to 808nm | 3-minute mean, positive response | Parameter~Wavelength+(1\|Subject) | 0.079972 | 0.009904 | 8.074717 | 171 | 0.909952 | 0.957764 | 0.060558 | 0.099382 |
| Wavelength; 1064nm compared to 808nm | 3-minute mean, negative response | Parameter~Wavelength+(1\|Subject) | 0.022664 | 0.010811 | 2.096383 | 175 | 0.059322 | 0.093252 | 0.001470 | 0.0438514 |
| Frequency; 40Hz compared to 10Hz | %BOLD online τ, positive response | Parameter~Frequency+(1\|Subject) | 5.538327 | 2.605602 | 2.125546 | 148 | 0.035681 | 0.035681 | 0.377121 | 10.730102 |
| Frequency; 40Hz compared to 10Hz | %BOLD online τ, negative response | Parameter~Frequency+(1\|Subject) | 5.323985 | 2.470107 | 2.155366 | 147 | 0.201989 | 0.485931 | 0.482575 | 10.165395 |
| Frequency; 40Hz compared to 10Hz | 3-minute mean, positive response | Parameter~Frequency+(1\|Subject) | 0.284527 | 0.077660 | 3.663752 | 171 | 0.197955 | 0.430798 | 0.132306 | 0.436734 |
| Frequency; 40Hz compared to 10Hz | 3-minute mean, negative response | Parameter~Frequency+(1\|Subject) | 0.039499 | 0.032787 | 1.204715 | 175 | 0.260135 | 0.314687 | -0.024773 | 0.103753 |
| Power; 7mw/cm2 compared to 5mw/cm2 | %BOLD online τ, positive response | Parameter~Power+(1\|Subject) | 6.452977 | 2.955788 | 2.183166 | 148 | 0.326643 | 0.368171 | 0.659633 | 12.246321 |
| Power; 7mw/cm2 compared to 5mw/cm2 | %BOLD online τ, negative response | Parameter~Power+(1\|Subject) | 5.140258 | 2.405758 | 2.136648 | 147 | 0.408456 | 0.835493 | 0.424972 | 9.855544 |
| Power; 7mw/cm2 compared to 5mw/cm2 | 3-minute mean, positive response | Parameter~Power+(1\|Subject) | 0.221522 | 0.091436 | 2.4227 | 171 | 0.326596 | 0.326596 | 0.042376 | 0.400664 |

| | | | | | | | | | | |
|---|---|---|---|---|---|---|---|---|---|---|
| Power; 7mw/cm2 compared to 5mw/cm2 | 3-minute mean, negative response | Parameter~Power+(1\|Subject) | 0.194757 | 0.071672 | 2.717337 | 175 | 0.421455 | 0.421455 | 0.054414 | 0.335086 |
| Power; 9mw/cm2 compared to 5mw/cm2 | %BOLD online τ, positive response | Parameter~Power+(1\|Subject) | 13.522095 | 6.313569 | 2.141751 | 148 | 0.401776 | 0.401776 | 1.147495 | 25.896685 |
| Power; 9mw/cm2 compared to 5mw/cm2 | %BOLD online τ, negative response | Parameter~Power+(1\|Subject) | 6.126335 | 2.771372 | 2.210578 | 147 | 0.140682 | 0.140682 | 0.694446 | 11.558224 |
| Power; 9mw/cm2 compared to 5mw/cm2 | 3-minute mean, positive response | Parameter~Power+(1\|Subject) | -0.00777 | 0.032233 | -0.241060 | 171 | 0.655610 | 0.655613 | -0.070947 | 0.055407 |
| Power; 9mw/cm2 compared to 5mw/cm2 | 3-minute mean, negative response | Parameter~Power+(1\|Subject) | 0.007923 | 0.049825 | 0.159017 | 148 | 0.311903 | 0.311903 | -0.089737 | 0.105577 |
| Nasal distance (mm) | %BOLD online τ, positive response | Parameter~Distance+(1\|Subject) | -1.149218 | 2.400432 | -0.47875 | 148 | 0.376243 | 0.376243 | -5.854065 | 3.555629 |
| Nasal distance (mm) | %BOLD online τ, negative response | Parameter~Distance+(1\|Subject) | 6.052357 | 2.777961 | 2.178705 | 147 | 0.304865 | 0.304865 | 0.607546 | 11.497154 |
| Nasal distance (mm) | 3-minute mean, positive response | Parameter~Distance+(1\|Subject) | 0.463494 | 0.032265 | 14.36523 | 171 | 0.535138 | 0.535138 | 0.400251 | 0.526729 |
| Nasal distance (mm) | 3-minute mean, negative response | Parameter~Distance+(1\|Subject) | 0.095218 | 0.079704 | 1.194645 | 148 | 0.268545 | 0.268545 | -0.061011 | 0.251432 |

### d) iPBM Linear Mixed Effects analysis results on offline (stimulus) effects

| Effect | Parameter | fitlme Formula | Estimate | SE | tStat | DOF | pValue | FDR_pValue | CI_Lower | CI_Upper |
|---|---|---|---|---|---|---|---|---|---|---|
| ITA | %BOLD offline τ, positive response | Parameter~ITA+(1\|Subject) | -9.581744 | 2.342022 | -4.091227 | 146 | 0.137023 | 0.203061 | -14.1721105 | -4.991381 |
| ITA | %BOLD offline τ, negative response | Parameter~ITA+(1\|Subject) | 6.065391 | 2.765593 | 2.193161 | 157 | 0.234154 | 0.289187 | 0.644829 | 11.485953 |
| ITA | 3-minute mean, positive response | Parameter~ITA+(1\|Subject) | -0.050158 | 0.042772 | -1.172683 | 173 | 0.043951 | 0.055424 | -0.133983 | 0.033683 |
| ITA | 3-minute mean, negative response | Parameter~ITA+(1\|Subject) | 0.146695 | 0.009706 | 15.113847 | 171 | 0.071337 | 0.079293 | 0.127666 | 0.165714 |
| Wavelength; 1064nm compared to 808nm | %BOLD offline τ, positive response | Parameter~Wavelength+(1\|Subject) | 6.097662 | 2.858929 | 2.132848 | 146 | 0.237447 | 0.286271 | 0.494161 | 11.701163 |
| Wavelength; 1064nm compared to 808nm | %BOLD offline τ, negative response | Parameter~Wavelength+(1\|Subject) | -1.161804 | 2.826633 | -0.411020 | 157 | 0.058692 | 0.089532 | -6.702005 | 4.378397 |
| Wavelength; 1064nm compared to 808nm | 3-minute mean, positive response | Parameter~Wavelength+(1\|Subject) | 0.141234 | 0.037877 | 3.728753 | 173 | 0.192441 | 0.249014 | 0.066991 | 0.215469 |
| Wavelength; 1064nm compared to 808nm | 3-minute mean, negative response | Parameter~Wavelength+(1\|Subject) | -0.037521 | 0.056662 | -0.662192 | 171 | 0.141305 | 0.209345 | -0.148578 | 0.073538 |
| Frequency; 40Hz compared to 10Hz | %BOLD offline τ, positive response | Parameter~Frequency+(1\|Subject) | 16.878114 | 5.918142 | 2.851927 | 146 | 0.005153 | 0.005153 | 5.155521 | 28.601041 |
| Frequency; 40Hz compared to 10Hz | %BOLD offline τ, negative response | Parameter~Frequency+(1\|Subject) | -4.467394 | 0.193191 | -23.124230 | 157 | 0.021890 | 0.054394 | -4.088740 | -4.846048 |
| Frequency; 40Hz compared to 10Hz | 3-minute mean, positive response | Parameter~Frequency+(1\|Subject) | 0.070297 | 0.033631 | 2.090244 | 173 | 0.028862 | 0.091171 | 0.136207 | 0.0043731 |
| Frequency; 40Hz compared to 10Hz | 3-minute mean, negative response | Parameter~Frequency+(1\|Subject) | -0.089862 | 0.003799 | -23.654120 | 171 | 0.390613 | 0.438032 | -0.082414 | -0.097306 |
| Power; 7mw/cm2 compared to 5mw/cm2 | %BOLD offline τ, positive response | Parameter~Power+(1\|Subject) | 2.961176 | 1.291041 | 2.293634 | 146 | 0.053957 | 0.065723 | 5.491616 | 0.430736 |

| | | | | | | | | | | |
|---|---|---|---|---|---|---|---|---|---|---|
| Power; 7mw/cm2 compared to 5mw/cm2 | %BOLD offline τ, negative response | Parameter~Power+(1\|Subject) | 2.800487 | 1.720493 | 1.627724 | 157 | 0.033494 | 0.054751 | -6.172653 | 0.571679 |
| Power; 7mw/cm2 compared to 5mw/cm2 | 3-minute mean, positive response | Parameter~Power+(1\|Subject) | 0.072980 | 0.006421 | 11.365831 | 173 | 0.015536 | 0.068782 | 0.060395 | 0.085565 |
| Power; 7mw/cm2 compared to 5mw/cm2 | 3-minute mean, negative response | Parameter~Power+(1\|Subject) | -0.013264 | 0.007944 | -1.669688 | 171 | 0.242952 | 0.307165 | -0.028832 | 0.002313 |
| Power; 9mw/cm2 compared to 5mw/cm2 | %BOLD offline τ, positive response | Parameter~Power+(1\|Subject) | 2.222573 | 0.981193 | 2.265174 | 146 | 0.080982 | 0.090221 | 0.299435 | 4.145711 |
| Power; 9mw/cm2 compared to 5mw/cm2 | %BOLD offline τ, negative response | Parameter~Power+(1\|Subject) | 4.112557 | 1.640098 | 2.507506 | 157 | 0.264988 | 0.264987 | 0.897965 | 7.327149 |
| Power; 9mw/cm2 compared to 5mw/cm2 | 3-minute mean, positive response | Parameter~Power+(1\|Subject) | 0.342388 | 0.023301 | 14.694133 | 173 | 0.320916 | 0.320948 | 0.2967152 | 0.388057 |
| Power; 9mw/cm2 compared to 5mw/cm2 | 3-minute mean, negative response | Parameter~Power+(1\|Subject) | -0.045961 | 0.029617 | -1.551845 | 171 | 0.077361 | 0.130942 | -0.104009 | 0.012089 |
| Nasal distance (mm) | %BOLD offline τ, positive response | Parameter~Distance+(1\|Subject) | 12.795082 | 5.989572 | 2.136226 | 146 | 0.779744 | 0.838296 | 1.055519 | 24.534641 |
| Nasal distance (mm) | %BOLD offline τ, negative response | Parameter~Distance+(1\|Subject) | 2.360452 | 1.033514 | 2.283909 | 157 | 0.055523 | 0.121380 | 0.334765 | 4.386139 |
| Nasal distance (mm) | 3-minute mean, positive response | Parameter~Distance+(1\|Subject) | -0.351945 | 0.032772 | -10.739200 | 173 | 0.083154 | 0.083015 | -0.416173 | -0.287707 |
| Nasal distance (mm) | 3-minute mean, negative response | Parameter~Distance+(1\|Subject) | 0.090353 | 0.044658 | 2.023220 | 171 | 0.122761 | 0.184503 | 0.002825 | 0.177882 |
